# Gravlax: an annotation-independent molecular evidence archive for single-cell RNA-seq

**DOI:** 10.64898/2026.09.18.752708

**Authors:** Rob Patro

## Abstract

A cell-by-gene count matrix is the artifact of a single-cell RNA-seq experiment that is most often stored, shared, and reanalyzed. It is the output of a computation whose inputs are the sequenced molecules and a gene annotation, and while the molecules never change, the annotation is revised continually. Once the matrix has been produced, the evidence behind it can no longer be reinterpreted. Recovering that evidence means returning to raw reads or alignments that are large, costly to process, and frequently unavailable.

We ask whether a concise representation of the molecules themselves can be extracted once and reused indefinitely, to quantify under any future annotation, to query and discover features that no annotation yet describes, and to pool evidence across cells and samples. Our starting observation is that the procedures that turn alignments into counts, gene assignment and UMI collapse, never read most of what an alignment file contains. They consume *relations* among molecules such as shared genomic geometry, shared placements, barcode identity, and the equality or near-equality of UMIs. We show that these relations form a statistic that is sufficient for such consumers, and we design a compact, seekable, content-authenticated archive that stores them while deferring every annotation-dependent decision to analysis time. Archives compose into content-addressed collections that route cohort queries to the molecules that can answer them without copying molecules. We implement these ideas in a tool called `gravlax`.

Across four human 10x 3′ datasets, `gravlax` archives require 11–18 bits per read and are 9.0–12.7× smaller than tag-preserving CRAM. Count matrices replayed from an archive deviate from direct `STARsolo` quantification by 0.24–0.75% of normalized UMI mass, whereas changing GENCODE v32 to v49 moves 2.12–4.64%, and quantification replay is 34– 82× faster than `STARsolo` at matched thread budgets. A federated index over eight archives occupies 2.96% of their size, answers a 96-query junction panel 2.59× faster than the archives alone, and screens the cohort genome-wide for unannotated splice events that recur across donors in just 9 seconds.

Because the molecules are retained, the archives also answer questions the matrix has discarded. An analysis of four peripheral-blood archives recovers a validated FYB1 immune-cell splicing switch, a cross-fitted fragment model appropriate for 3′ chemistry reveals an eight-donor shift in NTRK2 terminal-isoform usage from astrocyte and neural-stem-cell populations to mature neurons, and pooling evidence across cells within the context of an expectation-maximization algorithm recovers 75–98% of withheld multi-gene molecule labels. `Gravlax` is open source, implemented in Rust, licensed under the BSD 3-clause license, and available at https://github.com/COMBINE-lab/gravlax.

## 1 Introduction

The most commonly produced, shared, and reprocessed artifact of a single-cell RNA-seq experiment is the count matrix which encodes, for each cell and each annotated gene, the estimated number of distinct molecules observed. A count matrix is the output of a computation with two inputs, the sequenced molecules and a gene annotation, that age very differently. The molecules are fixed the day the library is sequenced. The annotation is revised several times a year. In a routine peripheral blood mononuclear cell (PBMC) dataset, moving from GENCODE^1^ v32 to v49 gives a 2.43% normalized matrix difference and changes the count of 21.8% of expressed genes by more than 10%; in brain nuclei the same change gives a 4.64% normalized matrix difference with intron-inclusive counting. Incomplete 3′ ends, intronic-read policies, and overlapping gene models can hide entire cell types and their markers^2^. In practice, the matrix is what is kept. Reads and alignments move to cold storage or are discarded, and the count matrix, having committed to one interpretation, becomes the object most downstream work builds on.

The natural remedy, reprocessing from raw reads, is rarely exercised. Even with fast quantifiers^3,4^, the reads for one experiment occupy tens of gigabytes and a cohort terabytes, a burden that many laboratories cannot absorb for every annotation release. What one might prefer instead is an intermediate object far smaller than the reads, but retaining what a future quantification would need, so that requantifying under a new annotation becomes a *query* rather than a *pipeline*. In this paper, we ask what such an object must contain, show that it can be made very small, and demonstrate that once it exists, it supports much more than requantification. We develop a tool, called `gravlax`, with a suite of operations to create, index, and query such annotation-independent archives.

Existing intermediate representations sit at the two ends of an uncomfortable tradeoff. Coverage resources such as recount3/Monorail and Snaptron^5,6^ support annotation-independent queries over bulk data, but their coverage-centric representation erases the cell-barcode and UMI relations on which single-cell deduplication depends. The BUS^7^ and RAD^4^ files produced by fast single-cell quantifiers retain those identifiers, and are generally much smaller than their BAM/CRAM counterparts, but they still bind each read’s mapping to the transcriptome chosen at mapping time, so a new annotation demands a new mapping. Reference-free systems such as sc-SPLASH^8^ and Malva^9^ support sequence-level discovery from cell-associated *k*-mer summaries, but do not preserve the joint barcode, UMI, genomic-placement, and splice-block structure that gene assignment consumes. At the other extreme, a tagged BAM or CRAM file is general enough to revisit any of these decisions, but pays for sequence, quality strings, read names, and per-read repetition that assignment procedures never examine, and is therefore large enough that it is reused far less often than the matrix derived from it. Boiler10 showed for bulk RNA-seq that a lossy alignment archive can still reproduce quantification. No comparable representation exists for single-cell data (i.e. a genome-coordinate, molecule-resolved statistic that leaves the annotation undecided). This is precisely the object we construct.

Our approach begins by examining the information used by gene assignment and UMI collapse. Assignment, as implemented in Cell Ranger^11^ and `STARsolo`^12,13^, inspects a placement’s aligned blocks, its splice junctions, its strand, and its alternative placements. UMI collapse inspects whether two UMIs in the same cell are equal or reside within a fixed distance bound. After alignment, neither inspects a read’s sequence, its qualities, or its name. This suggests storing the outcomes of those comparisons rather than the values themselves, i.e. an equivalence class in place of a UMI string, a shared shape in place of repeated block coordinates, and a shared pattern in place of repeated alternative placements. UMI collapse operates within a (cell, gene) context, and that context does not exist until an annotation is supplied. Collapsing molecules at construction time would therefore fix one of the decisions we want to revisit. The resolution is to store the *relations* (UMI equality and adjacency, shared geometry, shared placement, cell identity) and to postpone every annotation-dependent partition of those relations to query time. The resulting object is a *quotient* of the alignments under the equivalence induced by the downstream procedure, so that two inputs that the procedure cannot distinguish are represented only once.

A second design decision concerns cohort queries like “does this splice junction recur in every donor, and in which cells?” `Gravlax` supports such queries directly through federated indexes, which separate *evidence* from *routing*. Counts are computed from the archives themselves. A derived collection records the content digests of its member archives and stores routes, i.e. small structures that identify which archives and compressed chunks could contain matching molecules. These indexes can be rebuilt, extended, or specialized while the molecular evidence remains in the source archives.

This paper makes four contributions. (i) We formulate annotation replay as the evaluation of a fixed downstream procedure on a statistic computed before the annotation is known, characterize which relations that statistic must retain, and identify the single empirical reduction we apply beyond it. (ii) We design a seekable, content-authenticated archive format that factors repeated UMI, geometry, and multimapping relations, and stores them in independently decodable genomic chunks with coordinate, junction, and cell indexes. (iii) We give exact algorithms for replay, molecular queries by cell or cell group, and content-addressed federation, including a junction-by-shape index that answers junction predicates across a cohort while reading a small fraction of the source. (iv) We show that retaining molecular evidence across cells and samples enables inference that a matrix cannot support and give specific examples of protocol-aware terminal-isoform analysis and cross-cell recovery of ambiguous molecules. Formal statements, format specifications, complete algorithms, and secondary analyses appear in Supplementary Notes S1–S8.

## 2 Methods

### 2.1 Problem formulation

We want an archive that is small, can be queried and decoded selectively, and stores exactly the molecular evidence needed to answer our questions. Genome alignment and barcode correction are performed once, and annotation-dependent steps are deferred to query time.

Formally, let *A* denote a fixed set of genome placements (intervals or chains of intervals produced by annotation-free alignment), let *B* be a fixed corrected-barcode map, let *g* be a gene annotation on the same reference coordinates, and let *C* be a fixed gene-assignment and UMI-collapse procedure, such as the “Gene” counting rule of `STARsolo`. We refer to *C* as the *consumer* of the archive.

A conventional pipeline computes *C*(*A, B, g*) directly. `Gravlax` instead computes a statistic *T* (*A, B*) before *g* is known, and later evaluates *C*(*T* (*A, B*), *g*) for whatever *g* is supplied (Figure 1). We say this statistic is *sufficient* for *C* if

**Figure 1.**
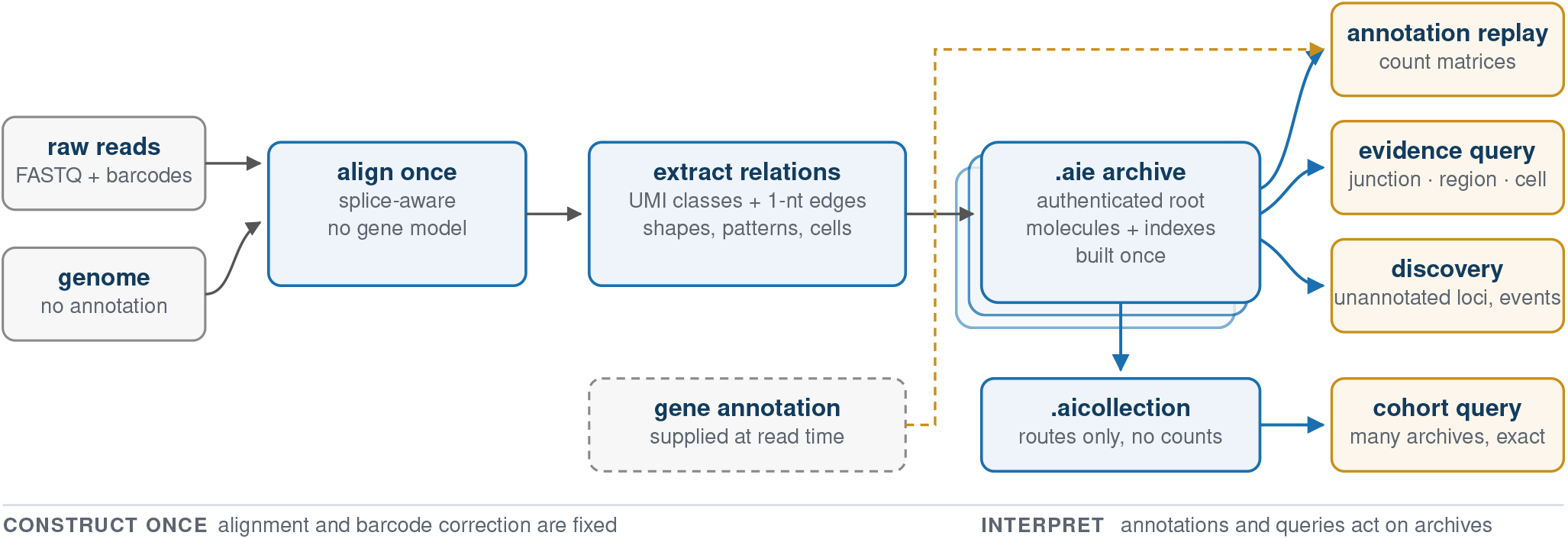
System overview. Alignment and barcode correction are performed once, without an annotation. The archive stores annotation-independent molecular relations and is the only authority for counts. An annotation supplied later drives replay, and derived collections route cohort queries back to the same evidence.

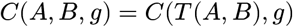

for every annotation *g* in the consumer’s domain. Sufficiency is thus relative to *C. T* must retain every field that *C* reads, and may omit everything else.

For the consumers we target, the fields read are the aligned blocks, junctions, strand, and alternative placements of each read, the identity of its cell, and the equality and one-mismatch adjacency relations among UMIs within a cell. Table 1 lists these relations alongside how `gravlax` stores them, and which decision about them is deferred. Each relation is also *necessary*, in the sense that deleting it produces two inputs that the archive cannot distinguish but that *C* maps to different outputs under some compatible annotation. Deleting a splice boundary, for instance, makes a spliced and a contiguous placement of the same span indistinguishable, yet an annotation whose exon boundaries match only the spliced placement counts them differently. Supplementary Note S1 gives such a counterexample for every relation. Analyses that also inspect bases or qualities (e.g. allele-specific or editing queries) require the original reads, or some alternative representation of the sequence information, which we do not currently store.

**Table 1.** What the archive stores and what it defers. Each row is a relation the downstream procedure observes. `Gravlax` stores it once and leaves every annotation-dependent partition to replay.

| observable relation | stored representation | decision deferred to read time |
| --- | --- | --- |
| UMI equality and adjacency | global class + per-cell graph | gene-restricted collapse |
| junction-chain geometry | shape + extremes + weight | transcript compatibility |
| alternative placements | anchor + shared differences | candidate-gene union |
| genomic locality | chunks + coordinate postings | which evidence to decode |
| archive identity | section digests + content root | collection binding |

`Gravlax`, by default, applies an additional, empirically evaluated reduction, *Q*. Within a UMI class, reads that share the same junction chain are replaced by their multiplicity and at most two coordinate-extreme representatives (i.e. the most extended placement and the most contained one). The intuition is that gene compatibility is decided by chain geometry and by where the chain begins and ends, so the reads in the interior of a deeply sequenced chain rarely change an assignment. We measure the incremental effect of *Q* by comparing compact and full-read (sans *Q* reduction) `gravlax` replay. Comparison with a direct annotation-aware `STARsolo` run also includes the effects of annotation-free alignment and differences in counting rules, giving the end-to-end fidelity reported below. Supplementary Note S1 separates these sources of error.

### 2.2 Storing relations instead of values

The dominant space cost of a read-level file is values that downstream computations never read, or read only to compare with other values. Of course, alignment itself needs the reads, but assignment and UMI deduplication then access a much smaller set of information. `Gravlax` stores that reduced information, and often the outcome of the comparisons, directly rather than the source data (Figure 2).

**Figure 2.**
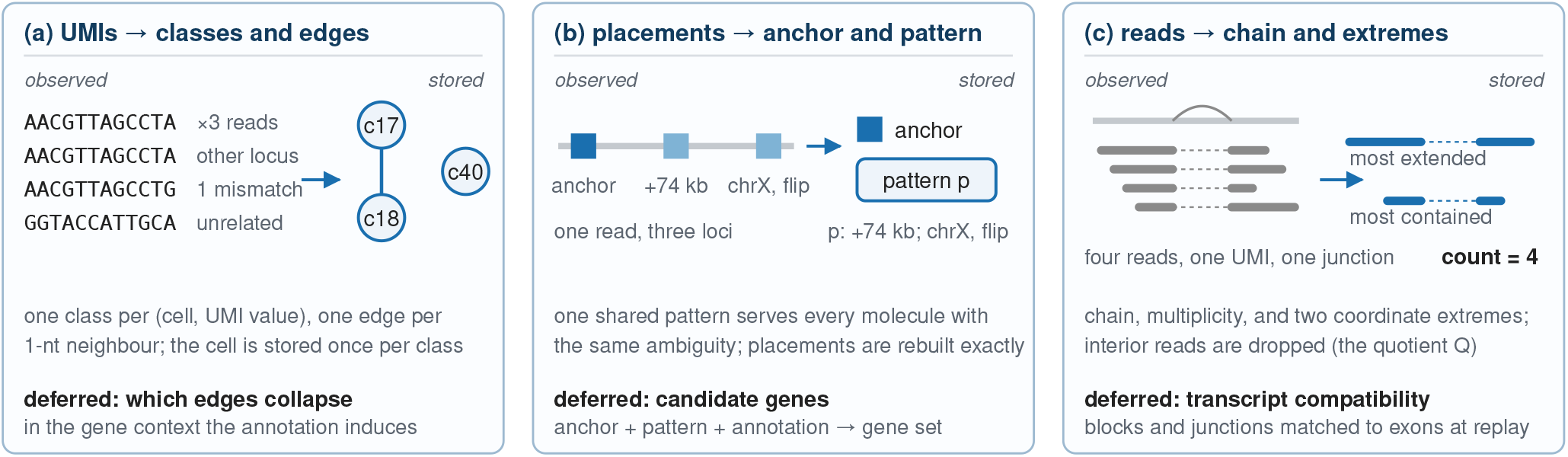
Relational factorization. Values are replaced by equality, adjacency, shared-placement, and shared-chain relations. Each replacement is either exact for the downstream procedure or, in the case of *Q*, has a separately measured empirical error.

#### UMIs become a graph

Every distinct (cell, UMI) pair receives a single global class identifier, even when the same pair is observed at several loci. Within a cell, observed UMI values at Hamming distance one are joined by an undirected edge. Because the graph is built over the whole cell, replay can later restrict it to whichever gene context the new annotation induces, so collapse remains deferred without any UMI string being stored. Cell identity is recorded once per class rather than once per molecule.

#### Geometry becomes a shape

A placement’s block structure, relative to its start, is stored once in a shared shape dictionary, so repeated splicing geometry is stored once and junctions are recovered from block boundaries rather than stored. Reads with the same chain share a shape and store a multiplicity plus their two extreme placements. A multimapper stores one anchor placement and a shared pattern of coordinate differences from that anchor describing its alternatives. The same pattern is referenced by every molecule that exhibits the same ambiguity.

#### Locality becomes an index

Molecules are sorted by anchor coordinate and partitioned into independently compressed genomic chunks. Within a chunk, each column is coded by whichever of delta and varint coding, rANS^14,^ absolute identifiers, or zstd^15^ compresses it best, chosen deterministically from the observed distribution. Separate range, junction, and cell postings map a query to the chunks that could satisfy it, so an operation selects chunks before decoding the information for any molecule. Class identifiers and graph edges are defined archive-wide. The cost of a query is proportional to the indexes and chunks it touches, and a full replay scans each chunk exactly once. The binary layout and compression methods are specified in Supplementary Notes S1 and S2.

### 2.3 Authenticated layout and replay

Selective decoding uses section-level content digests. Each `gravlax` archive (an `.aie` file, for annotation independent evidence) carries an authenticated directory whose entries name a canonical section, its offset and lengths, and the BLAKE3^16^ digest of its exact compressed payload (Figure 3). A content-root digest covers the header, the directory’s location, and the directory bytes, with a format-specific prefix. Opening an archive authenticates the directory; selecting a section authenticates its compressed bytes before bounded decompression. A second identity, computed over the ordered section names, lengths, and digests, is invariant to offsets and container wrapping, so it recognizes the same evidence after rewrapping or relocation. Together these establish that two files encode the same evidence.

**Figure 3.**
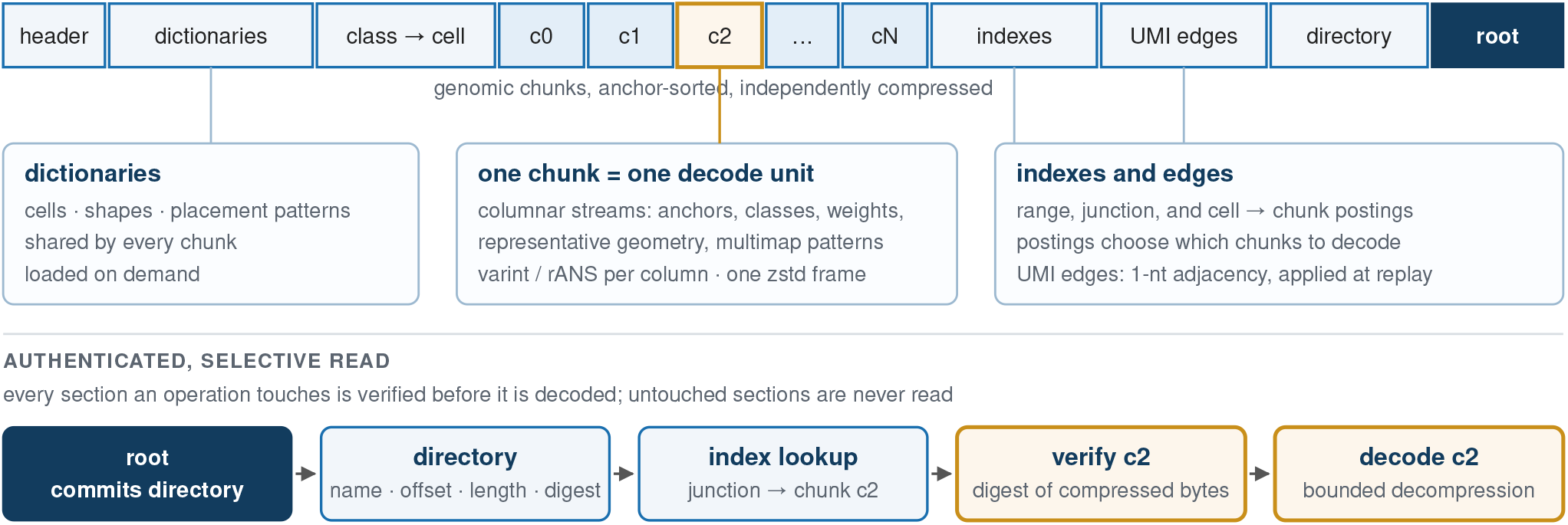
Seekable authenticated archive. The directory root covers every compressed section digest. Indexes select independent chunk decode units, and every section an operation consumes is authenticated before it is decoded.

Replay is where we make the decisions deferred during construction. A supplied GTF is compiled into transcript spans, merged exons, strands, and an overlap index. A placement is considered compatible with a transcript when all of its blocks are exonic, its junctions coincide with consecutive exon boundaries, and its strand obeys the library’s rule. This defines gene candidates, matching the Gene rule of `STARsolo` and Cell Ranger. `Gravlax` also implements Gene-Full, which overlaps aligned blocks with exon-derived gene spans, including introns, on the accepted strand. A skipped intron alone does not create an overlap. Candidate genes are unioned over primary and alternative placements. Uniquely assigned rows are grouped by (cell, UMI class), the `STARsolo MultiGeneUMI_CR` rule retains the gene with greatest read support, and within each (cell, gene) the classes are ordered by support and merged greedily through the one-mismatch graph17 restricted to that gene.

Replay decodes at most two chunks per worker, writes compact assignments into 64 cell-partitioned buckets, and collapses one bucket at a time, so peak memory is governed by the chunk window rather than by the dataset. Supplementary Notes S2 and S3 give the compatibility rules and the complete algorithm.

### 2.4 Content-addressed federation and junction-by-shape indexing

Beyond single-sample archives, `gravlax` supports federated queries across cohorts. A separate `.aicollection` index records each member archive’s content digest, offset-independent evidence identity, chromosome digest, genome signature, and location hint. An external manifest maps content identities to current paths, allowing archives to move without rebuilding the collection. New archives are added through index layers linked to their parent by its content root. Queries check source identities and combine entries across these layers.

Coordinate and junction catalogues identify which archives and chunks might contain a junction, but confirming that a candidate molecule carries a specific junction would still require loading each source’s full shape dictionary. The *junc tionbyshape index* removes this cost by exploiting a simple invariant. The genomic coordinates of a junction depend on where a placement starts, but the *intron span* (i.e. acceptor minus donor coordinates) is a property of the shape alone. For each pair of adjacent blocks in a shape *q*, let *o* be the donor offset within the shape and *l* the intron span. The collection construction enumerates the triple (*l, q, o*) for every such pair, sorts and deduplicates the (*q, o*) pairs under each *l*, and delta-codes them in independently compressed blocks of at most 256 spans. Construction uses one packed triple array and makes two passes over each source dictionary. For *m* adjacent-block pairs, sorting costs *O*(*m* log *m*) work and the encoded index is linear in the number of distinct triples.

A query for the junction (*d, a*) computes *l* = *a* − *d*, binary-searches for the block containing *l*, and obtains the candidate (*q, o*) pairs. A placement of shape *q* beginning at position *p* carries the requested junction exactly when *p* + *o* = *d* and *p* + *o* + *l* = *a*. The per-archive counting procedure checks both boundaries and then counts distinct UMI classes. The index includes every adjacent block pair in the source dictionary and records that dictionary’s digest and number of shapes. Lookup costs *O*(log *b* + *r*_*l*_ + *x*) for *b* index blocks, *r*_*l*_ candidate pairs under the span, and source work *x* selected by those candidates. Sources indexed only by coordinate (i.e. not participating in the junction-by-shape index) are queried through their own shape dictionaries. Supplementary Notes S2 and S3 give the index definitions and pseudocode.

### 2.5 Queries by cell and cell group, and event discovery

The `gravlax` query layer treats a UMI class, rather than a read, as the unit of evidence. To reduce comparisons, range and junction predicates first union their chunk postings, so a batched plan decodes each selected chunk once, and cell filtering is applied before any group or bulk reduction. For a splicing event, a class *c* contributes inclusion and exclusion indicators *I*(*c*) and *E*(*c*) where the conservative counts are

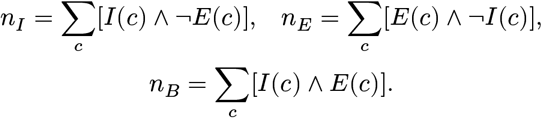

Every informative class falls in exactly one category, and a class that supports both alternatives contributes to the shared-support count *n*_*B*_.

Annotation-free discovery enumerates alternative donors, alternative acceptors, and three-junction cassettes directly from catalogue coordinates, without a gene model. An inverted component-to-event map turns all candidates into one packed reduction over sorted (event, class, mask) tuples. Across a cohort, coordinate keys are joined over genome-matched archives while preserving the distinction between an event that is absent from a sample’s catalogue and one that is present with zero support in a group. Minimum-support requirements are checked first in the sample or cell group with the lowest estimated query cost. Sparse output stores nonzero counts and defines omitted rows as zero. Supplementary Note S3 gives the algorithms and the composable GQ query language that can be used to frame and execute complex queries across `gravlax` archives. Supplementary Note S5 defines the biological comparisons.

### 2.6 Inference on retained evidence

Retained molecular evidence supports models of fragment geometry and ambiguous gene assignment that a count matrix cannot. We describe two examples in this section.

#### Protocol-aware terminal inference

Retaining fragment boundaries invites questions about 3′-end usage, but the measurement model must match the assay. In fragmented 10x 3′ libraries, the observed cDNA boundary lies a variable distance, often hundreds of bases, upstream of the true cleavage site. To model how 3′-end usage changes between cell types, we take PolyASite18 sites in unique terminal regions, filtered for internal priming, as candidates and estimate a kernel *K*(*d*) over the fragment-to-site distance from molecules compatible with exactly one site (without reference to cell-group labels). To keep the assay model independent of the biological contrast, donor *i* is fitted with *K*_−*i*_ learned from the other donors. If endpoint *e* is compatible with site *s*, its responsibility, normalized over the sites compatible with *e*, is

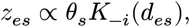

and an EM algorithm alternates responsibilities and site mass. Gene-level usage is the expected, UMI-weighted, transcript-oriented site rank, and its contrast between cell groups within each donor is the quantity tested across the cohort. Supplementary Note S6 gives eligibility rules, calibration, sensitivity analyses, and the definition of terminalsite groups.

## Crosscell recovery of ambiguous molecules

A single cell’s unambiguous evidence can sometimes identify the most likely gene among its candidates, but the abundance estimate is unstable when that cell’s coverage is sparse. Pooling unambiguous evidence across all cells supplies a more reliable prior, with the greatest gains where per-cell coverage is thinnest. For a multi-gene-only class *k* in cell *c* with candidate set *C*_*k*_, `gravlax` implements a pooled EM that assigns responsibility

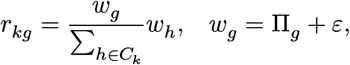

where *ε* = 10^−9^ and Π_*g*_ is the sample-wide abundance, initialized from, and repeatedly anchored to, unambiguous classes. Normalization restricts the comparison to genes that could have produced the molecule. The archive reconstructs *C*_*k*_ from its primary and alternative alignments. Responsibilities form a separate additive count layer. Supplementary Note S7 gives the estimator, a group-aware extension, and calibration analyses.

### 2.7 Evaluation design

We use a direct annotation-aware `STARsolo` run^12^ as the reference for end-to-end comparisons. Full-read versus co-ordinate-extreme `gravlax` replay isolates the effect of the approximation *Q*. GeneFull is the primary counting model for *D*_2′_ brain nuclei, while the other datasets use Gene. Matrix comparisons retain fixed called barcodes and match unversioned Ensembl identifiers. Normalized matrix deviation is half the *L*_1_ distance divided by reference UMIs. Storage comparisons distinguish tag-preserving CRAM19 from a sequence-free molecule CRAM that encodes exactly the relations the archive stores, in a generic container. Runtime comparisons use equal total thread budgets, with method order randomized and balanced across repetitions and warm-cache runs retain the operating system’s file cache.

Per-stream entropy accounting and the content-matched molecule CRAM quantify the storage contribution of the representation, and region queries provide a comparison for junction-specific indexing. Biological analyses use donor cross-fitting and label permutations. Masked gene recovery compares pooled, per-cell, and uniform estimators on the same candidate sets. Biological samples are the replicates for cohort inference. We report size as logical bytes, runtime as wall time, and memory as peak resident set. Supplementary Notes S4 and S8 provide details of the datasets, parameter choices, and hardware used.

### 3 Results

### 3.1 Does the archive preserve annotation replay?

The archive exists precisely so that the annotation can be changed after the fact, and so the first question is whether the error from deferring the annotation is small compared with the signal that motivates deferring it. We built one archive from the PBMC dataset *D*_0_ and replayed it, without rebuilding, under GENCODE v32, v48, v49, and a v49 variant with 900 expressed features withheld.

Relative to a direct annotation-aware `STARsolo` run under each annotation, full-read replay deviation is 0.207–0.246%, the compact archive deviates by 0.244% under v49, and restoring the withheld features by replay recovers 99.88% of the mass they had removed. Across the larger PBMC dataset *D*_1_, the glioblastoma dataset *D*_2_, and the brainnucleus dataset *D*_2′_, deviation is 0.307–0.746% (Table 2). The matched v32-to-v49 annotation difference is 2.12–4.64% (Figure 4, left), or 5–10× larger. For brain nuclei, GeneFull replay gives 71.16M collapsed UMIs versus 70.82M from `STARsolo` on the same 6,460 nuclei, and its 0.746% deviation is 6.2× smaller than the 4.64% annotation difference. We note that the archive itself is unchanged when switching from Gene to GeneFull. Supplementary Note S4 separates counting-model effects, nucleus calling, and whole-archive assignment statistics.

**Table 2.** Principal validation datasets. Deviation is half-*L*_1_ divided by matched `STARsolo` UMIs under GENCODE v49 (human) or M39 (mouse). Brain nuclei use GeneFull on fixed barcodes; other rows use Gene.

| dataset | reads | cells | archive | replay deviation |
| --- | --- | --- | --- | --- |
| $D_0$ PBMC | 66.6M | 1,225 | 111.1 MB | 0.24% |
| $D_1$ PBMC | 383.9M | 5,038 | 544.0 MB | 0.31% |
| $D_2$ glioblastoma | 250.7M | 5,573 | 439.7 MB | 0.43% |
| $D_{2'}$ brain nuclei | 263.4M | 6,460 | 595.2 MB | 0.75% |
| $D_5$ mouse 5' | 33.7M | 1,129 | 62.4 MB | 0.75% |

**Figure 4.**
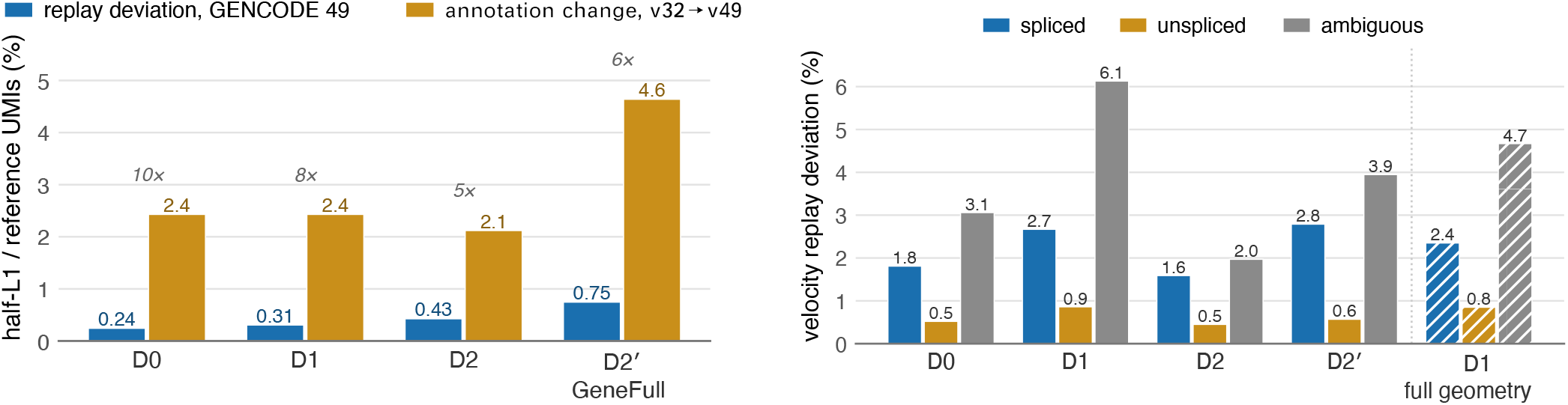
Replay fidelity and the effect of read reduction. **Left**, normalized matrix deviation from replay stays below that from changing the annotation; brain nuclei use GeneFull. **Right**, RNA-velocity component deviations under the two-representative reduction. Unspliced stays below 0.9% and the ambiguous component is most sensitive to deep chains. Hatched bars are *D*_1_ rebuilt with every unique-read geometry retained.

Of course, small matrix deviations could still affect biological conclusions. In a Gene-only sensitivity analysis, we tested cell clusters and their markers. For each dataset we selected the finest Leiden^20–22^ partition that was stable under subsampling and reseeding of the `STARsolo` matrix alone, then held those parameters fixed for the replayed matrix. The medoid partitions agree between methods with adjusted Rand index 0.965–0.997 and top-25 marker overlap 0.970–1.000 across four datasets, and between-method instability exceeds within-method instability by at most 0.024.

Fine-scale neighborhoods vary, but clusters that are stable under subsampling and reseeding of the oracle matrix are equally stable between the oracle and replay. The mouse 10x 5′ dataset (*D*_5_), processed with annotation-free two-pass alignment and the opposite strand rule, has 0.751% normalized matrix deviation. In fact, the counting rule matters far more than how the matrix is produced. On fixed brain nuclei, under a PCA/k-means cluster analysis, `STARsolo` versus replay GeneFull gives ARI 0.980, whereas Gene versus GeneFull under replay gives 0.333. Supplementary Note S4 separates this exploratory comparison from the Gene-only controls.

Finally, the cost of *Q* for Gene replay is small. On *D*_0_ under the v49 annotation, deviation is 0.24% with two coordinate extremes and 0.22% on the full-read replay. This 0.03-point difference measures the incremental effect of *Q*. The next section accounts for the rest.

### 3.2 What accounts for the remaining deviation?

Even once *Q* is accounted for, two sources of deviation remain, namely the read reduction itself (which Gene counting tolerates well, though other consumers may not) and the alignment.

RNA-velocity counting^23^ may be more sensitive to read reduction than Gene counting. It classifies a molecule as spliced, unspliced, or ambiguous by intersecting the transcript sets of all reads in a UMI class, and should expose whatever the interior reads of a chain carried. Across the four human datasets, the spliced, unspliced, and ambiguous matrices deviate from direct processing by 0.45–6.1% (Figure 4, right). The unspliced component, which drives most velocity inference, stays below 0.9%; the ambiguous component carries most of the discrepancy. With the `STARsolo`-derived neighborhood graph, PCA projection, and gene-wise unspliced-to-spliced ratios held fixed, the median per-cell velocity-vector cosine is 0.993–0.999 and the median Jaccard similarity of the top three transitions is 1.0.

Stratifying by evidence completeness reveals the underlying mechanism. Matrix entries whose chains retain every read agree with direct processing at the 1–2% level in every component. At the deepest saturation, *D*_1_, ambiguous entries with complete evidence deviate by 1.5%, compared with 6.5% for entries in which two representatives stand in for three or more reads. The discrepancy therefore increases with saturation. Retaining every distinct unique-read geometry (the optional geometry-fidelity encoding, Supplementary Note S2) tests this directly on *D*_1_: the archive grows from 11.3 to 14.4 bits per read, Gene deviation falls from 0.307% to 0.253%, and the spliced and ambiguous deviations fall from 2.67% and 6.13% to 2.35% and 4.67%, with unspliced unchanged at 0.85% (Figure 4, right, hatched). Read reduction therefore accounts for about a quarter of the ambiguous discrepancy. Per-read terminal signals are a separate case as a tail-bearing read may be an interior read removed by reduction, so the optional sparse tail representation records these events before reduction (Supplementary Notes S1 and S6).

The larger part of the remaining deviation arises in alignment itself. `Gravlax`’s ingest alignment is unseeded, whereas `STARsolo` aligns against an index seeded with the annotation’s splice junctions. Junctions present during STAR13 indexing can change the reported alignments, since mappings that cross annotated junctions receive a scoring bonus. Seeding the *D*_0_ ingest alignment with the junction list from GENCODE v32, five years older than the replay annotation, reduces the Gene deviation under v49 from 0.244% to 0.160%, close to the 0.143% obtained with v49 junctions, and reduces the unspliced velocity deviation from 0.52% to 0.04% and the ambiguous deviation from 3.06% to 1.51%. This works because junction sets are far more stable than transcript sets across releases (99.1% of v32 junctions persist in v49, and junctions added between v32 and v49 carry only 0.2–0.7% of junction reads in our datasets (Supplementary Note S4)). A seeded alignment therefore remains annotation-independent with respect to gene models, and the archive records the seed and its digest in its provenance. We provide this capability as a (non-default) option in `gravlax`.

### 3.3 Is the representation compact and fast?

Routine reuse requires compact storage and fast access, so we measure archive size and reanalysis time.

Across the four human datasets, archives occupy 111–595 MB, or 11–18 bits per read: 9.0–12.7× smaller than tag-preserving CRAM and 32–49× smaller than compressed FASTQ (Table 3). Tag-preserving CRAM includes sequence, qualities, and read names. For a content-matched comparison, we built a sequence-free molecule CRAM that stores the same placements, weights, UMI classes, molecule groups, and UMI graph as the archive, as generic records. `Gravlax` is still 2.10–2.55× smaller than this baseline, and because the baseline must expand 27–253 million generic records to answer any question, `Gravlax` enables quantification 6.64– 13.69× faster than the CRAM archives across the matched human and mouse evaluations.

**Table 3.** Storage ladder. FASTQ and tag-preserving CRAM retain richer read-level information. The molecule CRAM encodes the same relations as the archive but indexes none of them.

| artifact | $D_0$ | $D_1$ | $D_2$ | $D_{2'}$ |
| --- | --- | --- | --- | --- |
| FASTQ | 5.0 GB | 26.6 GB | 17.6 GB | 19.0 GB |
| tag-preserving CRAM 3.1 | 1.40 GB | 6.52 GB | 4.16 GB | 5.36 GB |
| content-matched molecule CRAM | 258.2 MB | 1.294 GB | — | 1.519 GB |
| **gravlax archive** | **111.1 MB** | **544.0 MB** | **439.7 MB** | **595.2 MB** |

Per-stream accounting for the native archive estimates less than 1% additional size reduction from replacing the entropy coder (under a memoryless value model). Thus, `gravlax`’s storage gains come from factoring repeated information: classes instead of UMI strings, shared patterns instead of repeated alternative placements, chains instead of reads, and cell identity per class rather than per molecule. The UMI relation costs 1.2 rather than 27 bits per molecule, and each alternative-placement pattern is shared by 7.9 molecules on average. Bits per molecule vary with saturation and composition, whereas bits per read stay within 11–18 (Supplementary Note S4).

Replay is fast enough to make requantification routine, since alignment is paid once at ingest and never repeated. At equal 24-thread budgets, Gene-only `STARsolo` with alignment output disabled takes 83.71, 472.90, and 326.43 s on *D*_0_, *D*_1_, and *D*_2′_ whereas Gene replay takes 2.44, 5.80, and 4.76 s, a 34–82× speedup. Under GeneFull on *D*_2′_, `STARsolo` takes 361 s and replay 9.3 s (39×) with 5.6× less peak memory. A 46.9MB compiled annotation replaces a 3.32GB GTF and reduces a fixed junction query from 1.19 to 0.09s. Storage normalization, timing protocols, and memory results are in Supplementary Note S4.

#### 3.4 What does a cohort query read, and what can it find?

We compared coordinate-only and junction-by-shape collection indexes over eight independently constructed archives from an adult human subependymal-zone (SEZ) cohort^24^ totaling 1.33 GB. The coordinate-only collection occupies 0.674% of the source and adding the junction-by-shape index brings it to 2.963%, with the additional index itself 0.885 times the size of the shape dictionaries it selectively replaces.

For fixed dense, sparse, and junction-set predicates, the junction-by-shape index reads 10.5%, 12.9%, and 10.5% as many source bytes as the coordinate-only collection respectively, runs 3.25–3.43× faster, and uses 4.53–5.29× less peak memory (Table 4). Across a 96-query panel of point junctions stratified by how many archives contain them, the median source-byte ratio is 0.119, the 95th percentile is 0.332, and aggregate wall time improves 2.59×.

**Table 4.** Junction-by-shape indexing relative to coordinate-only execution for fixed predicates; values below one are improvements. Region queries use the coordinate index.

| predicate | source bytes | wall time | peak memory |
| --- | --- | --- | --- |
| dense junction | .105 | .308 | .189 |
| sparse junction | .129 | .292 | .221 |
| junction set | .105 | .308 | .191 |
| region control | 1.000 | 1.038 | 1.000 |

Batch queries reuse the same index. For example, a 32-predicate plan with high locality takes 0.06 rather than 0.93s because chunk lists are combined before decoding.

Notably, discovery over the same collection needs no coordinates. A single command pools the junction catalogues of the eight archives (1.34 M junctions), enumerates every cassette whose components recur in at least two archives (2.98 M candidates), counts exact UMI-class support per archive and cell group, and keeps candidates seen in at least four donors with at least eight informative UMI classes and support in two required cell groups. Genome-wide, this takes 9 s and 3.0 GB of memory at 24 threads, reading the archives once. Of 205,735 candidates that no GENCODE v32 transcript explains, 3,442 contain a junction absent from the annotation, 2,408 of them in all eight donors, including the FNBP1 cassette of Section 3.5. Running discovery on each archive and merging instead takes 33 s and proposes only events whose components all occur in one archive’s catalogue, so it misses recurrences like FNBP1. Supplementary Notes S2–S4 describe index construction, the query comparisons, and the discovery protocol.

### 3.5 Can retained evidence answer questions the matrix cannot?

We present three analyses that use retained molecular evidence: a cohort splice-event comparison, a protocol-aware model of 3′-end usage, and a coordinate-free discovery of a recurrent cassette.

#### Splicing differences across PBMC samples

We compare T cells and monocytes in four PBMC experiments, defined using marker-gene rules on the STARsolo GENCODE v49 matrices. Candidate events and counts come from exact junction-set queries in each archive. Of 20,028 events recurring in the catalogues, 784 have at least ten informative molecules in both groups in every dataset with a concordant direction, and 44 shift by at least 0.10 everywhere. FYB1 leads the list: cassette inclusion is 5.7–8.8% in T cells and 98.3–99.1% in monocytes in every dataset (Figure 5), concordant with RT-PCR validation^25^. CD47 provides a second previously validated high-effect locus^26^. Both differences recur across the four archives.

**Figure 5.**
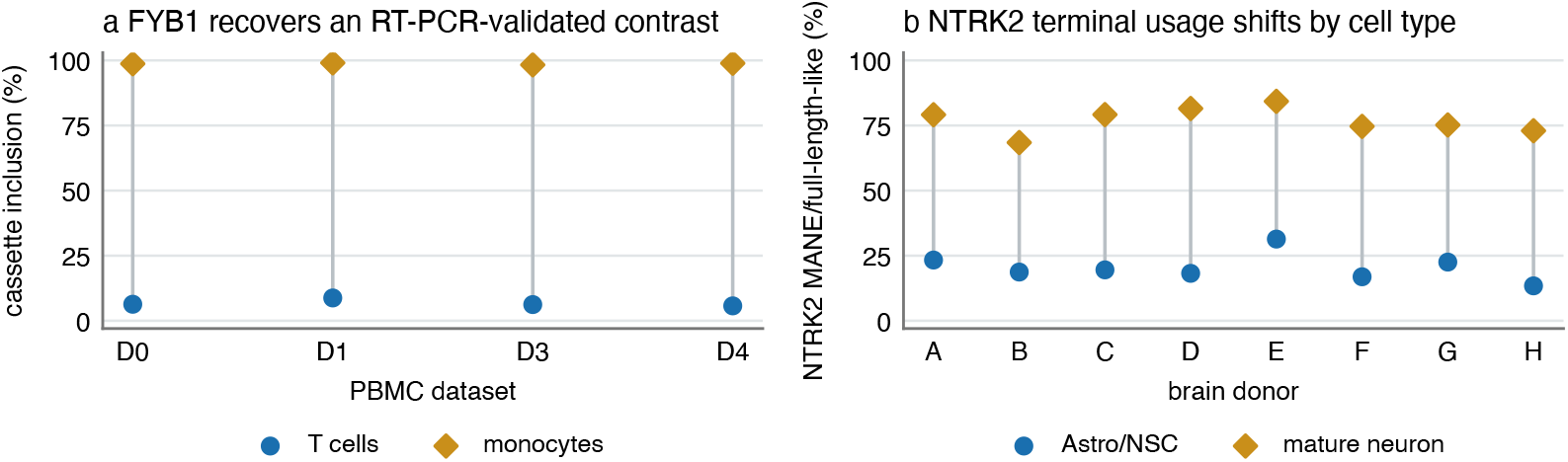
**a**, FYB1 cassette usage separates marker-defined T cells and monocytes in four PBMC datasets. **b**, cross-fitted terminal-site inference shifts NTRK2 toward the terminal group containing the full-length MANE Select transcript in mature neurons across all eight SEZ donors.

#### Terminal-site usage in 3′ libraries

The cross-fitted PolyASite mixture of Section 2.6 was fitted to 45,705 recurrent sites and 35.7M expected UMIs across the eight SEZ donors, contrasting astrocytes and neural stem cells (Astro/NSCs) with mature neurons. Held-out likelihood improves in every donor, 93 of 95 reported genes retain the direction of their usage difference under both neighboring site-merging choices, and a within-donor label shuffling reports no genes. Across all eligible genes the median shift toward distal sites is small (0.013 in distal usage index) though positive in every donor.

NTRK2, which encodes the TrkB receptor, is the strongest result (mean distal-usage effect +0.282, *q* = 1.48 × 10^−5^, eight of eight donor effects positive). Clustering GENCODE coding ends independently of the fit separates a proximal group of sites, associated with transcripts of median 1,431-nt coding sequence, from a distal group containing the 2,466-nt MANE Select^27^ full-length transcript. Usage of the distal group rises from 20.0% in Astro/NSCs to 78.1% in mature neurons (Figure 5; median paired change +56.8 percentage points, exact sign-flip *p* = 0.0078). This agrees with the known predominance of truncated TrkB.T1 in astrocytes and full-length TrkB in neurons^28^. We note that the analysis is exploratory, since model choice and NTRK2 selection used the same cohort. The inferred quantities are terminal-group usage fractions; their coding-end associations come from the annotated transcripts.

The genome-wide discovery scan of Section 3.4 surfaces a third example from the same SEZ cohort, this time drawing on splice geometry rather than fragment boundaries. FNBP1 encodes an F-BAR domain protein involved in membrane curvature and endocytic dynamics and is broadly expressed in the brain. An annotated 183-nt cassette in FNBP1 is supported by 44 UMI classes across 44 nuclei in six of eight donors and shows strikingly tissue-specific inclusion. We observe near-complete inclusion in brain-derived archives (94–100%) contrasted with low inclusion in PBMC archives (4–13%), a pattern corroborated in an independent normal brain dataset^29^. This recurrence is invisible to per-archive discovery, as the cassette’s three junction components are not jointly present in any single archive’s catalogue, so a per-archive merge sees evidence from at most two donors and rejects the event. Pooling junction catalogues across the full cohort before counting is what makes the cross-donor recurrence observable. Full junction co-occurrence patterns and cell-type localization are in Supplementary Notes S5.3 and S5.4.

The three analyses draw on different parts of the archive. Junction sets use observed splice geometry alone, both for the PBMC immune-cell comparison and for the FNBP1 discovery. The terminal-site mixtures combine archived fragment boundaries with a protocol model that could be learned only because molecules from many donors were retained. The FNBP1 result additionally illustrates that federated counting over a pooled catalogue can surface recurrent events that per-sample discovery would discard. Because the archive separates molecular evidence from its interpretation, the same file supports new annotations, genomic queries, and cohort comparisons as the questions change.

### 3.6 Does pooling across cells recover ambiguous molecules?

In *D*_0_, multi-gene-only UMI classes add a potential 6.3% over single-gene-supported classes and are dropped from conventional unique counting. Because the archive retains each such molecule’s candidate set together with the unambiguous evidence of every other cell, it can attempt to recover them. In a masked experiment, we took molecules whose unique evidence identifies their gene, removed that evidence down to the original candidate set, and asked each estimator to recover the hidden label. On fixed called barcodes, pooled EM attains 74.9–98.3% conditional top-1 recovery, compared with 68.9–96.6% for per cell EM and 43.2–57.9% for uniform allocation (Figure 6). For brain GeneFull, pooling raises recovery from 68.9% to 74.9% on 156,884 evaluable masked UMI classes. Another 74,467 of 231,351 masked classes (32.2%) lose their unique-evidence label from the remaining candidates and are excluded. We note that the counting model changes both candidate sets and the eligible truth population, so these scores measure recovery of withheld evidence, not biological ground truth.

**Figure 6.**
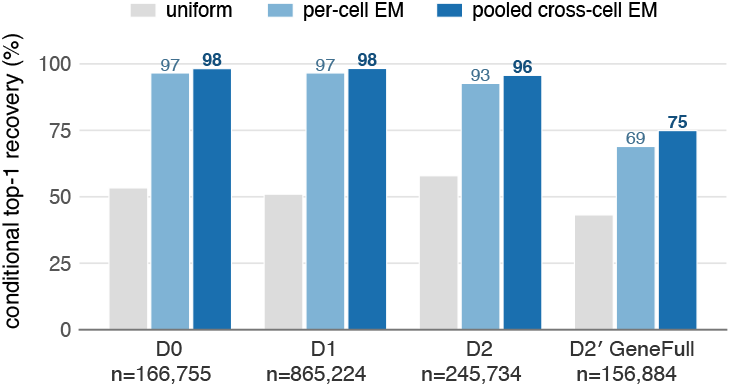
Conditional recovery of withheld unique-evidence labels on fixed called barcodes. Brain nuclei use GeneFull; other datasets use Gene. *n* counts evaluable masked UMI classes, before one-mismatch collapse.

Applied to *D*_0_ ambiguous classes, the estimator adds this inferred mass, 85% of it at an EM responsibility above 0.8. Compared to an independent pipeline, `alevin-fry`^4^, which performs its own fresh mapping from read sequence, 79% of the 254 most affected genes move closer, and recovered fractions agree between two PBMC datasets at *r* = 0.992.

The inferred counts form a separate layer alongside the replayed matrix. Their utility and calibration depend on the counting model and target population. Supplementary Note S7 reports calibration, alternative estimators, and the full protocol.

## 4 Discussion

`Gravlax` changes what the durable and sharable artifact of a single-cell experiment can be. A count matrix is cheap because it commits to one interpretation and discards the rest. A read archive is reusable because it retains far more than quantification consumes, but it is simply too large and burdensome for routine transfer and reanalysis, and can waste recomputation.

Between these extremes sits a reference-guided but annotation-free molecular evidence archive that stores relational statistics tailored to useful biological queries. It keeps equality and adjacency in place of UMI strings, shared shapes and patterns in place of repeated placements, and independently indexed chunks that serve replay and selective queries alike. Content-addressed collections extend the same object across samples while every count is still computed from the archives themselves. Thus, quantification can be replayed in seconds, a junction can be located across a cohort of independently constructed archives, a 3′ protocol can be given a new measurement model, and discovery can be run across a whole collection, all without returning to FASTQ or BAM/ CRAM files.

The alignment and the barcode correction remain fixed once an archive is built. The alignment may be seeded with a junction list, which the archive records, but it cannot be redone. Seeding is optional and off by default, and deserves deeper investigation since it can further reduce deviation from fresh runs. Read sequence and qualities are not retained, so analyses that use bases require the original reads.

What the archive discards after alignment is governed by *Q*, whose cost we have measured directly. That cost is very small for gene counting, and larger but still reasonable for counts that depend on every read in a UMI class. Retaining every unique-read geometry recovers about a quarter of that larger cost for 27% more storage on *D*_1_, and sparse terminaltail records preserve per-read tail events that reduction would otherwise remove.

Federated queries provide sample-level observations for cohort analyses. Statistical comparisons use biological replicates and an experimental design appropriate to the question, as in the paired donor contrasts presented here. If molecular evidence can be kept compactly and faithfully as we show here, and such representations are adopted, then the lasting artifact of a single-cell experiment need no longer be a frozen matrix of counts, but instead can be an archive of the molecular evidence itself. With an appropriate toolset, such archives can be requantified, requeried, explored and pooled across laboratories for as long as the biology they hold is worth revisiting.

### 4.1 Disclosure

R.P. is a co-founder of Ocean Genomics Inc. Claude and Codex were used in the development of this work and manuscript.

## 4.2 Acknowledgements

This work was supported by the US National Institutes of Health (R01HG009937) and by grants 2022-311195 and 2024-342821 from the Chan Zuckerberg Initiative DAF, an advised fund of the Chan Zuckerberg Initiative Foundation.

