## Supplementary Material for "Gravlax: an annotation-independent molecular evidence archive for single-cell RNA-seq"

**Rob Patro**

These notes describe the formal sufficiency argument and its counterexamples (Note S1), the archive and collection formats and their integrity model (Note S2), complete replay, query, and federation algorithms (Note S3), extended fidelity, storage, and runtime protocols (Note S4), biological analyses and their controls (Note S5), terminal-site analyses (Note S6), multimapper inference (Note S7), and datasets, software, and computational methods (Note S8).

#### Contents


S1 Evidence sufficiency and error from read reduction ... 3
S1.1 Fixed alignments and counting rules ... 3
S1.2 Necessity counterexamples ... 3
S1.3 Reduction of reads sharing a junction chain ... 3
S1.4 Effect of read reduction on RNA velocity ... 4
S2 Archive and collection formats ... 4
S2.1 Evidence construction ... 4
S2.2 Optional evidence and access representations ... 5
S2.3 Archive layout and content digests ... 5
S2.4 Collection indexes and archive relocation ... 5
S2.5 Indexing junctions by alignment shape ... 6
S3 Replay, query, and federation algorithms ... 6
S3.1 Barcode correction at construction ... 6
S3.2 Annotation compilation and compatibility ... 6
S3.3 Gene assignment and UMI collapse ... 7
S3.4 Two-pass construction of the junction-by-shape index ... 7
S3.5 Junction lookup through the collection index ... 7
S3.6 Event counts by cell and cell group ... 7
S3.7 GQ: a composable language for archived evidence ... 8
S3.7.1 Query structure and units of observation ... 8
S3.7.2 Predicates, quantifiers, and results ... 8
S3.8 Event discovery and the packed multi-event reducer ... 8
S3.9 Molecular splice graphs and replicate-aware inference ... 9
S3.10 Concurrent execution, streaming, and sparse results ... 9
S4 Fidelity, storage, and runtime evaluation ... 10
S4.1 Fidelity metrics and annotation replay ... 10
S4.2 Brain nuclei: counting model, calling, and units ... 10
S4.3 Annotation stability and junction-seeded alignment ... 11
S4.4 Mouse replay ... 11
S4.5 Gene-only stability of cell clusters and expression summaries ... 12
S4.6 Capability-matched storage ... 12
S4.7 Runtime and memory protocols ... 13
S4.8 Effect of the junction-by-shape index on query cost ... 14
S4.9 Cohort-wide event discovery over the collection ... 14
S5 Biological query and discovery analyses ... 14
S5.1 Splice-event differences between T cells and monocytes ... 14

S5.2 Observed junction components at AKR1A1 ... 15
S5.3 FNBP1 exon inclusion across tissues and cell groups ... 15
S5.4 FNBP1 junction co-occurrence in SEZ donors ... 16
S5.5 Terminal fragment-boundary redistribution ... 16
S5.6 Recovery of genes added in a later annotation ... 16
S5.7 Reference optimization as replay ... 17
S6 Terminal-boundary, PolyASite, and tail analyses ... 17
S6.1 Fragment geometry and analysis design ... 17
S6.2 Candidate sites and assay geometry ... 17
S6.3 Predictive and resolution controls ... 18
S6.4 NTRK2 terminal groups ... 18
S6.5 Non-templated poly(A)-tail evidence ... 18
S7 Inference for multi-gene evidence ... 19
S7.1 Evidence classes and pooled EM ... 19
S7.2 Masked recovery and leakage controls ... 20
S7.3 Candidate-normalized partial pooling ... 21
S7.4 Depth-dependent combination ... 22
S7.5 Leakage-controlled group construction ... 22
S7.6 Parameter selection and uncertainty ... 23
S7.7 Memory use ... 23
S8 Datasets, software, and reproducibility ... 23
S8.1 Evaluation datasets ... 23
S8.2 Genome and annotation identity ... 24
S8.3 Compute and software environment ... 24
S8.4 Timing and memory measurement ... 24
S8.5 Software and archived workflows ... 24
References ... 25

### S1 Evidence sufficiency and error from read reduction

#### S1.1 Fixed alignments and counting rules

Let $A$ be a set of genome placements after alignment, $B$ a corrected-barcode map, $g$ a gene annotation on the same reference coordinates, and $C$ a fixed assignment-and-collapse algorithm. A placement comprises its half-open aligned blocks, junctions implied by adjacent blocks, strand, alternative placements, cell identity, UMI relation, and read weight. The archive is constructed as $T(A, B)$ without observing $g$. A *compatible* annotation uses the recorded contig names and lengths and falls within the transcript-compatibility and UMI policy implemented by $C$.

The conditional sufficiency statement is

$$C(A, B, g) = C(T(A, B), g)$$

for every compatible $g$, provided $T$ retains every field that $C$ reads. It requires fixed alignments, barcode correction, and counting rules. An algorithm that uses sequence, base quality, read names, or a different molecule-collapse relation requires a statistic retaining those inputs.

#### S1.2 Necessity counterexamples

Each retained relation is necessary for the named policy in the usual pair-of-inputs sense. The following pairs agree after one relation is removed but yield different outputs under a compatible annotation.

| removed relation | indistinguishable inputs | distinguishing reading |
|---|---|---|
| strand | otherwise equal sense and antisense placements | a stranded transcript annotation |
| splice boundary | junction-crossing and contiguous placements with the same span | exons whose consecutive boundaries match only the spliced placement |
| alternative placement | unique and ambiguous evidence at the anchor | a second compatible gene at the alternative locus |
| cell identity | the same molecule geometry in two cells | the per-cell count matrix |
| UMI equality | two observations of one value and two distinct values | distinct-molecule counting |
| UMI adjacency | neighboring UMI values with equal geometry | directional one-mismatch collapse |

Table S1: Pairs of inputs establishing the necessity of each retained relation for the specified counting rule.

After alignment and barcode correction, this counting rule uses only alignment geometry and molecule relationships. Of course, allele, editing, and sequence-search analyses also require the original bases or qualities.

#### S1.3 Reduction of reads sharing a junction chain

We denote the default compact archive by $Q(T)$. For reads with one junction chain, $Q$ retains multiplicity, the most-extended placement (smallest start, largest end as a tie break), and the most-contained placement (largest start, smallest end as a tie break). If there are at most two distinct reads, all are retained. Gene compatibility is, in practice, usually determined by chain geometry and coordinate extremes. The effect of this reduction depends on the annotation and the counting rule, as quantified below.

Two comparisons distinguish the effect of read reduction from end-to-end differences in quantification:

- Coordinate-extreme and full-read Gravlax differ by 0.03 percentage points of moved UMI mass on $D_0$/GENCODE v49 (0.24% versus 0.22%). This measures the incremental effect of $Q$ for Gene replay.
- A direct annotation-aware STARsolo run also differs in junction discovery, placement, tie-breaking, and other counting rules. Comparison with STARsolo measures their combined effect on quantification.

The UMI graph preserves equality and one-mismatch adjacency as relations. The encoded relation costs 1.2 rather than 27 bits/molecule for repeated UMI values. Alternative alignments share patterns of coordinate differences from their primary alignment, with 7.9× mean reuse. On fixed alignments, grouping differs by 0.23% from STARsolo's `MultiGeneUMI_CR`/1MM behavior. This deviation simply reflects different collapse rules, since the conditional equality in Section S1.1 requires the same counting rules.

##### S1.4 Effect of read reduction on RNA velocity

RNA-velocity components depend on evidence from every read in a UMI class. Across the four human datasets, component deviation is 0.45–6.1%. Unspliced deviation remains below 0.9%, while the ambiguous component is most sensitive to deep chains. Entries whose chains retain every read agree with direct processing at the 1–2% level in every component. On $D_1$, ambiguous entries with complete evidence deviate by 1.5%, compared with 6.5% for entries in which two representatives stand in for at least three reads. To measure how much of the deviation the reduction itself causes, we rebuilt $D_1$ with the optional `--geometry-fidelity` mode described in Note S2, which retains every distinct accepted unique-read geometry with its multiplicity, and replayed both archives with the same binary (version 0.2.2). The fidelity archive occupies 692.7 MB (14.44 bits per read) against 544.0 MB (11.34 bits per read) for the compact archive, a 27% increase. Construction took 7.4 min at 24 threads. Gene deviation on the 5,038 called cells falls from 0.307% to 0.253%. Velocity component deviations fall from 2.67% to 2.35% (spliced), 0.86% to 0.85% (unspliced), and 6.13% to 4.67% (ambiguous). With every chain complete, the completeness stratification collapses to a single stratum. The two-representative reduction therefore explains roughly a quarter of the ambiguous-component deviation on the most saturated dataset, and the remainder arises upstream of the archive, in annotation-aware alignment and transcript-set classification.

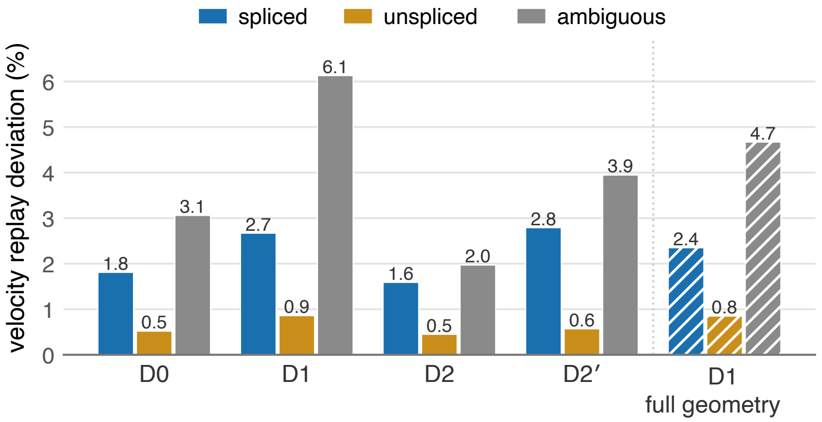


Figure S1: Velocity-component deviation by dataset. The ambiguous component is most sensitive to replacing deep chains with two coordinate extremes.

Read-specific terminal signals provide another counterexample to exact read reduction. The only signal-positive read may lie between the retained extremes, and one reduced molecule can summarize several distinct terminal anchors. Preserving these signals requires a list of their read-specific terminal coordinates. The optional terminal-tail representation extracts this list before reducing read geometry (Note S6).

#### S2 Archive and collection formats

##### S2.1 Evidence construction

Inputs are coordinate-sorted genomic alignments produced without a gene annotation. Barcode correction is performed during construction and fixed thereafter. Each primary alignment is normalized to 0-based half-open blocks, strand, junction chain, alternatives, corrected cell, UMI class, and read weight. Reference sequence is joined by contig name. Read sequence, quality, and name are not retained by the replay statistic.

Identical (cell, UMI) values receive a global class identifier across loci. An alignment's *shape* is its ordered list of aligned blocks expressed relative to its genomic start. Repeated shapes share one dictionary entry. Within each cell, observed UMI values at Hamming distance one define an undirected edge. By default, chain-equal reads are reduced as described in Note S1. A multimapping read stores its primary alignment and a shared pattern encoding the differences between primary and alternative alignments. Records sorted by genomic start are partitioned into independently compressed chunks. Per-stream delta/varint, rANS, or zstd coding is selected deterministically. Range-directory, junction-catalogue, and cell-to-chunk indexes are separate sections loaded on demand.

Classes and graph edges are defined across the archive. A class whose evidence crosses a physical chunk boundary retains one cell identity and one set of graph relations.

##### S2.2 Optional evidence and access representations

The optional `--geometry-fidelity` setting stores each distinct accepted unique-read geometry once with its exact multiplicity, within the original cell/UMI/locus record. This replaces the coordinate-extreme reduction for those reads. The additional geometry can affect gene assignment and velocity counts. Archive metadata records the reduction rule used during construction.

The `--access-index` option adds lists of chunks containing each repeated UMI class, fixed genomic bin overlapped by an aligned block, or exact junction. Separate lists cover uniquely mapped reads, unique reads plus primary alignments of multimapping reads, and all retained alignments including alternatives. These lists select candidate chunks. Decoded evidence determines matches. The `--chunk-records` option targets smaller chunks within genomic bins while keeping equal-anchor records together. Both options trade additional indexing or framing space for selective access. The `--compression-tuning` option instead selects lossless cell-map and shape-dictionary encodings by final compressed size, including run-length cell maps and shared splice/internal-block skeletons with per-shape terminal geometry. Shape identifiers and decoded evidence are unchanged by this compression option.

We note that the storage and fidelity results below describe the default compact archive.

##### S2.3 Archive layout and content digests

The archive begins with magic header (`AIE0`) and typed sections. Each section has a canonical name, physical offset, raw length, compressed length, and BLAKE3 digest of the exact compressed payload. The authenticated directory contains those entries. A content root is a BLAKE3 digest calculated from the header, directory location, and directory bytes, with a format-specific prefix.

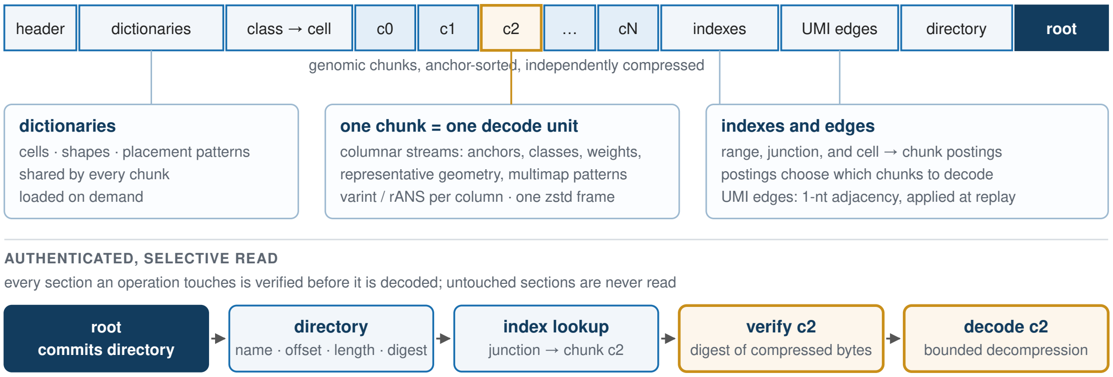


Figure S2: Default authenticated archive layout. The content root commits the section directory and compressed payload digests. Selected sections are hashed before bounded decoding.

Opening an archive verifies the directory root, canonical layout, and inline section headers without reading molecule payloads. Selecting a section hashes its compressed bytes before decompression and enforces the declared raw length. A full-verification option visits every section.

A second digest identifies the encoded evidence from ordered section names, raw and compressed lengths, and compressed-payload digests. It remains the same when section offsets change and changes with the encoded section contents or order.

Alignment provenance is bound to the archive root and records input identities, construction parameters, and the reference genome. Checking the directory root and section digests verifies that the bytes read match the recorded content identity.

##### S2.4 Collection indexes and archive relocation

The collection (`.aicollection`) is a separate index that locates evidence across sample archives. Its source manifest records the native archive root, offset-independent encoded-evidence identity, chromosome digest, and genome signature. Paths are merely location hints, since source identity depends on the encoded content. Construction reads source directories, metadata, and indexes. Each sample identifier and encoded evidence source occurs once in the collection.

A separate location manifest maps content identities to current local paths. Relative paths are resolved against the manifest's directory, allowing a bundle to be moved intact. Expected identities remain those recorded in the authenticated collection. The manifest can locate both archives and parent collection layers without modifying their bytes or rebuilding indexes.

Junction entries are partitioned into 16-Mb bins by splice-donor coordinate. Each entry records which samples contain the junction, an upper bound on support, and the chunks to inspect. New samples can be added as a separate index layer linked to the existing collection by its content root. Queries combine entries across these layers.

Opening a source checks its directory against the archive root and evidence identity recorded in the collection. Selected sections are checked against their digests as they are read. Full-content verification reads every section. Relocation requires only a location-manifest lookup in addition to these ordinary reads. The existing index remains usable wherever the matching archive content is located.

##### S2.5 Indexing junctions by alignment shape

For adjacent blocks $(a_i, l_i)$ and $(a_{i+1}, l_{i+1})$ in shape $q$, define $o_i = a_i + l_i$ and $\ell_i = a_{i+1} - o_i$. The builder emits packed triples $(\ell_i, q, o_i)$ in two passes over the shape dictionary. It sorts and deduplicates the triples, partitions them into blocks of at most 256 distinct spans, and delta-codes the $(q, o_i)$ pairs. The index records the source root, shape-section digest, number of shapes, and intron-span range of each block. Construction stores the triples in one array and writes compressed blocks sequentially.

At query time, the junction catalogue first selects sources and molecule chunks. The index block for the requested intron span supplies candidate shape/offset pairs. Each candidate placement is counted only when its reconstructed donor and acceptor coordinates equal the query. Upper bounds on support allow early exclusion of candidates below a requested count threshold.

#### S3 Replay, query, and federation algorithms

##### S3.1 Barcode correction at construction

Barcode correction fixes cell identities at archive construction. The input carries the raw 16-nt barcode and its Phred+33 qualities in `CR` and `CY` tags and is matched to a fixed whitelist. A first annotation-free pass over nonsecondary, nonsupplementary records counts exact whitelist hits. In the second pass an exact hit is accepted immediately. A barcode with more than one non-ACGT base is rejected. A barcode with one `N` enumerates the four bases at that position. Otherwise all whitelist neighbors at Hamming distance one are enumerated.

For candidate whitelist barcode $b$ differing at position $i$, its weight is

$$(f_b + 1)10^{-\frac{\min(Q_i, 33)}{10}},$$

where $f_b$ is the first-pass exact-hit count. Missing quality uses error probability 0.01. For a single `N`, the substitution is treated as a free mismatch and the weight is $f_b + 1$. The maximum-weight candidate is accepted only when its share of total candidate weight is strictly greater than 0.975. Otherwise the record is unassigned. The corrected barcode is then propagated to all placements of that read. This reproduces the `1MM_multi_Nbase_pseudocounts` decision without consulting an annotation.

##### S3.2 Annotation compilation and compatibility

Only GTF exon records with gene, transcript, and strand are consumed. Overlapping or duplicate exons within a transcript are merged. The optional compiled annotation stores normalized gene and transcript dictionaries, exons, strands, transcript spans, and an overlap index in a deterministic checksummed format.

A placement is compatible with a transcript when it lies in the transcript span, every aligned block is exonic, every junction matches consecutive exon boundaries, and the alignment/transcript strands satisfy the library rule. A gene is a candidate when any transcript is compatible. Candidate genes from alternative placements are unioned. Ordinary Gene counting retains only singleton candidate sets. GeneFull instead uses the exon-derived outer span of each gene, separately by chromosome and strand, and accepts any aligned block overlapping that span. Intronic blocks count, though skipped introns alone do not, and junction concordance is not required. Candidates are unioned over placements before the same unique-assignment and UMI rules.

##### S3.3 Gene assignment and UMI collapse

Classified rows are grouped by (cell, UMI class). `MultiGeneUMI_CR` retains the gene with greatest read support, breaking ties by deterministic gene order. Within each (cell, gene), classes are ordered by support and class identifier. Directional collapse[1] greedily merges through archived one-mismatch edges. Surviving roots form integer matrix entries. STARsolo, on the other hand, uses lexical UMI order in corresponding ties, and this policy difference contributes to the end-to-end deviation from STARsolo.

Bounded replay decodes at most two chunks per worker and partitions assignment records into 64 groups of cells for separate processing. Each group is sorted and collapsed independently. Eager replay performs the same classification and collapse with a larger active record set.

##### S3.4 Two-pass construction of the junction-by-shape index

Index construction reads a source shape section twice, so that its only large temporary allocation is a single contiguous, sortable triple array. The first pass counts adjacent-block pairs. The second fills an array of that size.

```
verify source root and compressed shapes digest
(n_shapes, m) <- count shapes and adjacent-block pairs
allocate an array of m triples
VISIT(shapes): emit (span, shape_id, donor_offset)
sort triples lexicographically
partition consecutive spans into blocks of at most 256 spans
delta-code sorted (shape_id, donor_offset) pairs in each block
record source root, shapes digest, and n_shapes; write index blocks
```

Algorithm S1: Two-pass construction of the junction-by-shape index. `VISIT` defines donor offset as the exclusive end of the left shape block and span as the next block's start minus that offset.

##### S3.5 Junction lookup through the collection index

The collection index selects candidate source archives and chunks. Counts are then computed from records in those chunks. For sources with a junction-by-shape index, lookup uses the matching shape entries. For other sources, it searches the source's shape dictionary.

```
1  authenticate collection chain and every referenced source identity
2  use coordinate catalogue to choose candidate sources and chunks
3  span <- query.acceptor - query.donor
4  if the source has no junction-by-shape index: search its shape dictionary
5  else binary-search the block interval containing span
6  authenticate/decode that block and obtain (shape_id, donor_offset)
7  for each candidate alignment, reconstruct donor and acceptor coordinates
8  retain the placement only if both coordinates equal the query
9  combine evidence within each UMI class; select requested cells or groups
10 emit the computed UMI-class counts for each sample
```

Algorithm S2: Junction lookup using the collection index to select source records for counting. Line 4 handles sources indexed only by genomic coordinate.

##### S3.6 Event counts by cell and cell group

Counts in the queries below are distinct UMI classes within one cell. A predicate is a condition evaluated on archived evidence. Range predicates use chunk intervals. Junction predicates use catalogue postings and, when present, the junction-by-shape index. A plan unions the chunk lists for all predicates before decoding, then OR-reduces within-class evidence.

Batch queries combine region and junction predicates. Execution takes the union of their postings, decodes every selected chunk once, and evaluates each molecule against the relevant predicates. Results retain the input query order. A junction absent from the catalogue is reported with `present=false` and zero counts.

For event $j$ and class $c$, let $I_{j(c)}$ and $E_{j(c)}$ indicate inclusion and exclusion evidence. Counts are

$$n_I = \sum_{c\left[I_{j(c)} \wedge \neg E_{j(c)}\right]}, \quad n_E = \sum_{c\left[E_{j(c)} \wedge \neg I_{j(c)}\right]}, \quad n_B = \sum_{c\left[I_{j(c)} \wedge E_{j(c)}\right]}.$$

Every informative class belongs to exactly one of these categories, so $n_I + n_E + n_B$ equals the number of classes with either bit. Usage is $\frac{n_I}{n_I + n_E}$ and shared support is reported separately. Cell filtering precedes group and bulk reduction, which ensures that group totals are sums over the same cell-level partition.

#### S3.7 GQ: a composable language for archived evidence

GQ expresses questions that combine genomic intervals, splice patterns, read support, and sample or cell metadata. A query specifies the observations to compare and how to summarize them. Named predicates and functions allow the same biological definition to be reused across archives and cohorts.

##### *S3.7.1 Query structure and units of observation*

A query starts with `from` and combines operations with the pipeline operator `|>`. `within` selects a genomic region, `where` filters observations, `derive` names calculated quantities, and `tally` or `summarize` aggregates them. `select`, `sort`, and `take` control the reported columns, ordering, and number of rows. `let` names reusable expressions. Nonrecursive `fn` definitions parameterize predicates such as minimum exon overlap or a junction pattern.

The source declares the counting unit. `.records` operates on retained cell/UMI/locus records, `.classes` combines records sharing an exact UMI value within a cell, and `.cells` combines a cell's records. `count()` counts the selected units, while `reads.total` and its unique and multimapping components report read multiplicities. Thus the source determines whether an aggregation counts records, UMI classes, or cells. Each cell and class is identified within its sample when several archives are queried together.

`within` selects units with a retained alignment overlapping the requested region. Subsequent class- or cell-level predicates inspect all records belonging to those selected units, including records outside that region. This allows a local genomic feature to define the population for a broader question about the same UMI classes or cells.

##### *S3.7.2 Predicates, quantifiers, and results*

Region literals (`g[...]`) are 0-based, half-open. Junction literals (`j[...]`) are exact aligned-block boundaries. Predicates test block overlap with a minimum number of bases, alignment starts and ends, junctions, and ordered junction paths, and annotation bindings expose gene spans, exon unions, and transcript junctions. Quantifiers state which observations must satisfy a predicate: `any unique { A & B }` requires one uniquely mapped alignment satisfying both, `any unique { A } & any unique { B }` allows different alignments of the same record, and `any record { ... }` lifts a record-level condition to a UMI class or cell. Predicates evaluate to true, false, or unknown, where unknown means the stored evidence does not resolve the question, for example an overlap that could lie on an omitted interior read; `tally` keeps all three states and `where` reports the unknowns it drops. Queries run unchanged over one archive or a federation manifest, and every result carries the query, plan, and archive digests. The following query tabulates exon-overlap and splice-junction support at PTPRC, including their co-occurrence on the same alignment:

```
header { gq = 1, assembly = "GRCh38" }
let locus = g[1:198600000..198800000]
let RA = overlaps(g[1:198696711..198696909:-], min: 12bp)
let RO = j[1:198692373..198703297:-]
from @pbmc.records
|> within locus
|> derive {
     ra = any unique { RA },
     ro = any unique { RO },
     joint = any unique { RA & RO }
   }
|> tally {ra, ro, joint} by {sample}
```

The complete language reference is at combine-lab.github.io/gravlax/cli/gq.

#### S3.8 Event discovery and the packed multi-event reducer

Coordinates are 0-based and half-open. An archived junction $(d, a)$ has donor $d$, the exclusive end of the left genomic block, and acceptor $a$, the start of the next block, with $d < a$. For two catalogue junctions satisfying the component-support threshold, an alternative-acceptor event has common donor and acceptors $a_0 < a_1$: $(d, a_0)$ is the coordinate-defined inclusion side and $(d, a_1)$ the exclusion side. An alternative-donor event has donors $d_0 < d_1$ and a common acceptor: $(d_0, a)$ is inclusion and $(d_1, a)$ exclusion. These labels follow genomic coordinate order. A cassette is the triple $[(d_0, a_0), (d_1, a_1); (d_0, a_1)]$

with $d_0 < a_0 \leq d_1 < a_1$: the first two flanks are inclusion and the long junction is exclusion. Candidates require observed support for every component. A GTF provides optional gene and strand labels after candidate generation and counting.

Component membership is inverted to event identifiers. One packed hit is `(event << 34) | (class << 2) | mask` in a `u64`: 30 event bits, 32 global UMI-class bits, and two side bits (`01` inclusion, `10` exclusion, `11` both). For each selected chunk, sorting and bitwise-OR reduction combine repeated representatives, alternative placements, and components before any group total is updated.

```
enumerate events from observed junction coordinates
invert each junction component to (event_id, side_bit)
union all component postings to obtain unique selected chunks
for each chunk in parallel: decode once
  for each molecule: inspect chain representatives and anchor multimappers
  OR matched side bits per event; emit packed(event,class,mask)
  sort by (event,class) and OR duplicate masks
merge chunk streams, sort, and OR again across chunk boundaries
map class to cell; filter requested cells before aggregation
stream event order, updating bulk or group I-only/E-only/both totals
```

Algorithm S3: Packed multi-event reduction. Grouping hits by event allows cell-group totals to be accumulated directly.

The packed organization avoids one hash table per event and decodes a selected chunk once even when it serves many predicates.

Federation unions event coordinate keys across genome-matched archives. The schema records `present=false` when a local catalogue lacks the structure, distinguishing absence from a present event with zero support in one group. A minimum-support condition requires $n_I + n_E \geq m$ in every requested sample or cell group. These groups are evaluated in increasing estimated cost so failure at a shallow archive or group prevents later decoding. Caller sample order is restored in the output.

##### S3.9 Molecular splice graphs and replicate-aware inference

For a locus, the catalogue selects junction coordinates and the union of their postings. Each selected chunk is decoded once. For each UMI class and strand, the algorithm forms the set of junctions observed among its retained reads, counting the class once. We call this strand-specific junction set a *molecular path fragment*, as it describes the splice junctions jointly supported by that UMI class. Multimapping reads contribute their stored primary alignment.

Path UMI and cell counts are formed first. Edge counts are then obtained by summing the retained path-fragment counts containing that edge. Thus, for every edge and every requested group, its reported UMI count equals the sum of the corresponding path counts by construction. Forward and reverse graphs remain separate and their edges are directed in transcript order. Catalogue support used to nominate an edge remains explicitly strand-combined.

Across samples, an edge enters the common graph when its local catalogue support reaches `min-support` in at least `min-edge-samples` archives. Every archive is then reduced against that common coordinate graph, including observed evidence below the discovery threshold. Missing observations are explicit zeros. Design rows must name unique sample identifiers and unique archive paths. An optional cell list restricts a sample before reduction.

For eligible sample $i$ and path $j$, let $k_{ij}$ be the path-fragment count and $n_i$ the count of all retained paths on the same strand. The model is $k_{ij} \sim \mathrm{BetaBinomial}(n_i, \mu\kappa, (1-\mu)\kappa)$. The null fits one mean and concentration. The alternative fits one mean per condition and one shared concentration. Deterministic profile likelihood restricts means to $[10^{-8}, 1 - 10^{-8}]$ and concentration to $[10^{-3}, 10^{6}]$. The likelihood-ratio statistic is compared with its one-degree-of-freedom asymptotic null, and BH correction covers every tested path in the locus. Testing requires the minimum number of biological replicates in each condition, sufficient path UMIs and supporting samples, and a same-strand comparator. Low-depth samples contribute descriptive counts. Each biological sample is one replicate.

##### S3.10 Concurrent execution, streaming, and sparse results

Chromosome workers share a fixed 24-thread budget. The summarizer visits one event object at a time and retains only qualified or selected records.

Sparse cohort output separates immutable metadata, the event catalogue, sample/event presence, and nonzero scoped counts. Omitted count rows are zero, allowing reconstruction of the dense sample-by-event result. Counts are reported separately for catalogue-recurrent candidates, candidates meeting minimum-support requirements, and candidates meeting the effect criterion.

On two whole-genome event analyses, applying minimum-support filters before counting retains 4,703/22,953 and 10,760/54,328 catalogue-recurrent events, reduces raw intermediate JSON from 413 to 102 MB, and completes in 45.3 s. Streaming summarization reduces peak RSS from 807 to 32 MiB. For 543 retained events, the sparse representation reconstructs all 3,258 sample observations from 76,970 bytes rather than 3,484,594 bytes of dense JSON, a 45.3× reduction. We note that the associated timing comparison is a single run and is reported descriptively.

### S4 Fidelity, storage, and runtime evaluation

#### S4.1 Fidelity metrics and annotation replay

We use a direct annotation-aware STARsolo run as the reference for end-to-end comparisons. Comparing full-read and compact `gravlax` isolates the effect of the two-representative quotient. Matrices are matched on fixed called barcodes and unversioned Ensembl identifiers, retaining distinct PAR-Y accessions. GeneFull is the primary quantification rule for $D_{2'}$ brain nuclei while other datasets use the Gene rule. Normalized matrix deviation is half the $L_1$ distance divided by the matched STARsolo UMI total. Annotation contrasts use STARsolo v49 as the denominator.

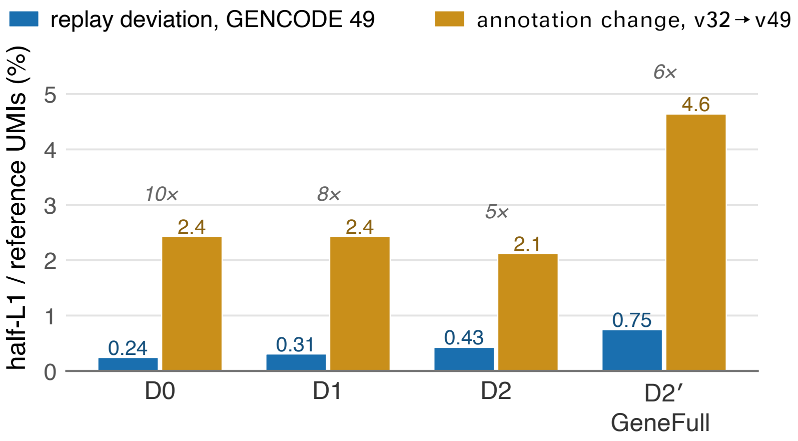


Figure S3: Replay deviation compared with the v32-to-v49 annotation signal. The former remains below the annotation change the archive is intended to recover.

One $D_0$ archive is replayed under v32, v48, v49, and a 900-feature withheld panel without reconstruction. End-to-end full-read deviation is 0.207–0.246% (0.244% for the compact archive under v49), and replay recovers 99.88% of the mass missing from the withheld matrix. Deviation is 0.307–0.746% on $D_1$, $D_2$, and $D_{2'}$. The matched STARsolo v32/v49 annotation difference is 2.43% on PBMC and 4.64% on brain nuclei. For GeneFull, the brain v49/v32 matrices contain 70,815,492/68,671,764 collapsed UMIs on the fixed nuclei and have an $L_1$ difference of 6,570,870 counts. Replay deviation is 0.746% at v49 and 0.733% at v32.

#### S4.2 Brain nuclei: counting model, calling, and units

All primary brain contrasts retain the same 6,460 Gene-called nuclei. Under v49, replay Gene gives 24,153,768 collapsed UMIs (median 2,882.5 UMIs and 1,833 detected genes per nucleus) while GeneFull gives 71,156,828 (median 8,390.5 and 3,532). Matched STARsolo totals are 24,280,804 and 70,815,492. GeneFull adds 48,631,181 and removes 1,628,121 counts across gene-by-nucleus entries, a net 47,003,060-count gain. We note that this counting-model effect uses the same archive, and is distinct from discovering new molecules or establishing more accurate biological assignments.

Independent STARsolo 2.7.11b EmptyDrops_CR[2] calling yields 6,460 Gene and 6,564 GeneFull nuclei, sharing 6,445 barcodes. Replay yields 6,458 and 6,565, sharing 6,446. GeneFull replay retains all 6,564 reference GeneFull calls and adds one. These independently selected cohorts do not, however, replace the fixed primary comparison.

A shared exploratory downstream analysis uses library normalization to 10,000, log1p counts, 2,000 genes ranked by pooled variance, gene z-scores, 30 joint PCs, and pooled k-means with 15 clusters (seed 2718) across the four matched matrices. Adjusted Rand index is 0.986 for STARsolo/replay Gene and 0.980 for GeneFull, but 0.333 for replay Gene/GeneFull. Thus, counting semantics affect clustering far more than replay in this analysis, though we note that the clusters are not cell-type ground truth. The separate PBMC splicing and eight-donor SEZ terminal-site analyses use their original inputs.

Whole-archive assignment statistics have different units from these matrices:

| unit | denominator | assigned Gene | assigned GeneFull |
|---|---|---|---|
| archive records | 130,270,701 | 30,672,329 | 81,557,405 |
| representative rows | 184,680,609 | 44,325,139 | 116,961,267 |
| UMI classes | 122,483,959 | 29,149,820 | 78,953,348 |

A record or class is assigned here when it contains at least one uniquely assigned representative. Correct record fractions are 23.55% and 62.61%. Multiple records can share a UMI class. Neither unit counts independently identified physical molecules. Full-input collapsed UMI totals are 28,630,629 for Gene and 77,643,934 for GeneFull, including uncalled barcodes.

##### S4.3 Annotation stability and junction-seeded alignment

Between GENCODE v32 and v49 the transcript set changed far more than the splice-junction set. Of v49′s 507,365 transcripts, 44% existed in v32 by identifier and 42% by exact intron chain. Of its 620,090 junctions, 61% existed in v32, and 99.1% of v32′s junctions persist in v49 (3,270 removed). Weighted by observation, the difference is smaller still. In the annotation-free two-pass alignments of $D_0$, $D_1$, and $D_{2'}$, junctions with at least three uniquely mapped reads that v49 contains and v32 lacks carry 0.24%, 0.32%, and 0.73% of junction reads, and junctions v32 contains and v49 lacks carry at most 0.03%. Junctions absent from every release carry 4.3%, 6.0%, and 15.7%. Transcript-level change over the same interval moves 2.4–4.6% of UMI mass through gene assignment.

The remaining difference between replay and a direct STARsolo run is therefore expected to arise largely from alignment, since STARsolo aligns against an index seeded with the annotation's junctions. To test this, we repeated the $D_0$ ingest alignment on the annotation-free index with a junction list inserted at mapping time (`--sjdbFileChrStartEnd`, overhang 90), using either the v32 or the v49 junction set, in two-pass mode (seed plus per-library discovery) and in one-pass mode (seed only). We archived each alignment with the seed file, discovery mode, and pass-1 catalogue recorded in provenance, and replayed it under v49 and v32 against the matching STARsolo run on the same 1,225 cells.

| ingest alignment | Gene v49 | Gene v32 | spliced | unspliced | ambiguous | archive |
|---|---|---|---|---|---|---|
| unseeded, two-pass | 0.244% | 0.232% | 1.81% | 0.52% | 3.06% | 111.1 MB |
| v32 seed, two-pass | 0.228% | 0.214% | 1.75% | 0.51% | 2.99% | 112.3 MB |
| v49 seed, two-pass | 0.213% | 0.223% | 1.58% | 0.50% | 2.65% | 112.5 MB |
| v32 seed, one-pass | 0.160% | 0.140% | 1.32% | 0.04% | 2.05% | 110.4 MB |
| v49 seed, one-pass | 0.143% | 0.154% | 1.09% | 0.03% | 1.51% | 111.5 MB |

Table S2: $D_0$ replay deviation from STARsolo by ingest alignment. Gene columns are moved UMI mass under the named annotation. Velocity columns are per-component relative $L_1$ under v49.

A v32 seed recovers most of the improvement a v49 seed gives, in both alignment modes. The one-pass seeded alignments agree with STARsolo more closely than the two-pass ones because the reference is itself a one-pass run on an annotated index, so junctions discovered per library are ones the reference never uses. We note that this measures agreement with the standard pipeline, not alignment accuracy. Archive size is unchanged by seeding.

##### S4.4 Mouse replay

For the mouse 10x 5′ v2 dataset, replay uses the opposite-strand relationship between alignments and transcripts and annotation-free two-pass alignment. Moved UMI mass relative to a GRCm39/GENCODE M39 STARsolo quantification is 0.751%.

| dataset | reads | cells | archive | bits/read |
|---|---|---|---|---|
| $D_0$ | 66.6M | 1,225 | 111.1 MB | 13.3 |
| $D_1$ | 383.9M | 5,038 | 544.0 MB | 11.3 |
| $D_2$ | 250.7M | 5,573 | 439.7 MB | 14.0 |
| $D_{2'}$ | 263.4M | 6,460 | 595.2 MB | 18.1 |
| $D_5$ | 33.7M | 1,129 | 62.4 MB | 14.8 |

Table S3: Archive sizes and read-normalized storage for the principal replay evaluations.

#### S4.5 Gene-only stability of cell clusters and expression summaries

These partition-selection controls retain the Gene counting model for all four datasets, including brain nuclei. The separate GeneFull sensitivity analysis is above. Clustering parameters are selected from the STARsolo matrices and then applied to replayed matrices. Across 15–40 principal components (PCs) and Leiden resolutions 0.10–0.80, we compare repeated clustering initializations, six 80% cell subsamples, and neighboring parameter choices. Agreement is measured by adjusted Rand index (ARI). The medoid is the partition with greatest mean ARI to the other partitions at the same parameter setting.

We select the finest partition with ARI at least 0.90 across initializations, 0.80 across cell subsamples, and 0.85 between medoids at neighboring settings. It must contain four to fifteen groups, each with at least 20 cells and at most 80% of the population. Marker reproducibility is measured by splitting cells into two deterministic halves, each with at least ten cells inside and outside every cluster, ranking genes by their mean expression difference between each cluster and the remaining cells, and comparing the 25 highest-ranked genes in each half. Their Jaccard similarity is the size of the intersection divided by the size of the union. The median similarity across clusters must be at least 0.50. The median Pearson correlation of gene-wise expression differences between halves must be at least 0.90.

| data | PC/res. | groups | within O/R | ARI/NMI | excess | markers |
|---|---|---|---|---|---|---|
| $D_0$ | 40/.60 | 7 | .995/.995 | .992/.988 | .003 | 1.000 |
| $D_1$ | 30/.40 | 9 | .951/.959 | .978/.970 | .008 | .991 |
| $D_{2'}$ | 40/.25 | 9 | .946/.925 | .965/.929 | .024 | .989 |
| $D_5$ | 20/.35 | 4 | .944/.944 | .997/.990 | -.001 | .970 |

Table S4: Stable macro-partitions under fixed STARsolo-selected parameters. Column definitions match the main paper.

Broad cell clusters are stable between STARsolo and replay, with medoid ARI 0.965–0.997. Fine-scale neighborhoods vary: median Jaccard similarity between each cell's two sets of 15 nearest neighbors is 0.765 on $D_1$ and $D_{2'}$. This local variation accompanies the sensitivity of clustering to initialization, subsampling, and resolution measured above. Replacing 20% of cell profiles with profiles from another cluster provides a perturbed-data comparison. Replay has 0.325–0.373 higher ensemble ARI and 0.186–0.296 higher median-neighbor overlap than this control. A steady-state velocity comparison holds the STARsolo-derived neighborhood graph, PCA projection, and gene-wise unspliced-to-spliced ratios fixed. It gives median cell-vector cosine 0.993–0.999 and top-three transition Jaccard 1.0.

The Spearman correlation between STARsolo and replayed pseudobulk profiles is 0.9989 on $D_1$, and the mouse $D_5$ Pearson and Spearman correlations are 0.9967 and 0.9980. For mouse 5′ velocity, component deviations are 2.91% spliced, 0.94% unspliced, and 4.62% ambiguous. The fixed-reference velocity comparison on $D_5$ gives median cell-vector cosine 0.9931 and top-three transition Jaccard 1.0. These values measure the stability of velocity vectors and neighboring-cell transitions under the shared STARsolo-derived model.

#### S4.6 Capability-matched storage

FASTQ and tag-preserving CRAM keep sequence, qualities, read names, and all alignments. Ingest-equivalent CRAM also retains barcode-correction inputs. The content-matched molecule CRAM contains sequence-free post-correction records plus documented tags for placements, weights, UMI classes, molecule groups, and the UMI graph. Gravlax interprets these molecule relationships during replay and supplies the genomic indexes used for selective queries.

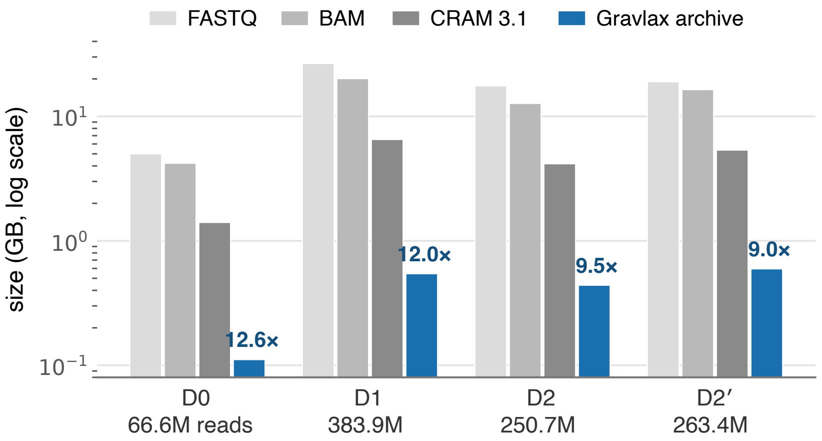


Figure S4: Storage ladder across the four human datasets. Ratios above archive bars are relative to tag-preserving CRAM 3.1.

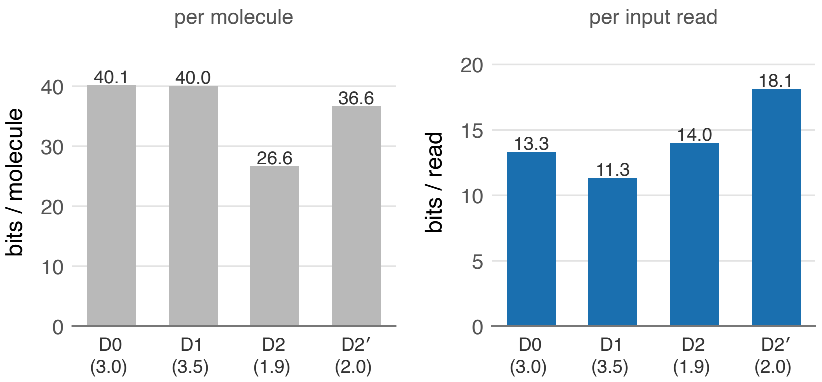


Figure S5: Normalization by reads and molecules. Bits/molecule reflects saturation and composition. The parenthetical labels give reads/molecule.

The archive is 9.0–12.7× smaller than tag-preserving CRAM and 32–49× smaller than FASTQ. Against the content-matched molecule CRAM it is 2.10–2.55× smaller.

Factoring UMI equality and one-mismatch adjacency requires 1.2 rather than 27 bits/molecule, while an alternative-alignment coordinate-difference pattern is reused 7.9× on average. On $D_0$/GENCODE v49, full-read Gravlax moves 0.22% of UMI mass and the stored two-extreme quotient moves 0.25%. The 0.03-point difference is the incremental error resulting from $Q$ for this Gene replay.

##### S4.7 Runtime and memory protocols

Runtime comparisons repeat both methods with equal total thread budgets, randomizing their order and balancing which method runs first. Warm-cache measurements retain the operating system's file cache between runs. Gene-only STARsolo disables BAM output. Molecule CRAM is decoded by `samtools` directly into the importer while both processes share one 24-CPU affinity set. No BAM is materialized. Sizes are logical bytes.

Five-run warm medians for STARsolo versus eager replay are 83.71/2.44 s on $D_0$, 472.90/5.80 s on $D_1$, and 326.43/4.76 s on $D_{2'}$, or 34.3–81.5× faster. These comparisons use the Gene counting rule. A matching GeneFull comparison on $D_{2'}$ (STARsolo with only GeneFull counting requested and alignment output disabled, versus `replay-rows --gene-full`; one advisory-cold observation per method, then five warm blocks in a balanced alternating crossover on the same 24 CPUs) gives warm medians of 361.5 s and 32.6 GiB for STARsolo and 9.27 s and 5.8 GiB for replay, a 39.0× speedup. All twelve runs reproduced their reference matrices exactly.

For cold-cache measurements, each input was evicted from cache and both methods incurred nonzero physical I/O. On $D_0$, STARsolo/replay takes 85.82/2.75 s (31.2×). The corresponding $D_1$ and $D_{2'}$ speedups are 58.4× and 61.6×. We note that these measurements characterize cold-cache behavior only for those inputs, under the recorded eviction procedure, on a shared host.

Streamed molecule-CRAM medians versus native replay are 20.65/2.34 s on $D_0$, 97.02/7.13 s on $D_1$, 86.19/6.29 s on $D_{2'}$, and 11.89/1.79 s on $D_5$. Native replay is therefore 6.64–13.69× faster.

For `gravlax` event listing, a deterministic 46.9-MB compiled annotation replaces the 3.32-GB v49 GTF. On a fixed annotated junction query, it reduces wall time from 1.19 to 0.09 s and memory from 376 to 136 MiB. Indexed region and junction queries return per-cell UMI counts in 0.05–0.3 s. On a $D_0$ interval, catalogue-only listing takes 0.01 s and exact cell counts take 0.06 s.

##### S4.8 Effect of the junction-by-shape index on query cost

The eight SEZ archives contain 1,326,036,691 bytes. A collection indexed only by genomic coordinate occupies 8,931,406 bytes (0.674% of the archive size). Adding the junction-by-shape index increases this to 39,286,118 bytes (2.963%). Construction takes 1.34 s and 607,132 KiB peak RSS, compared with 0.705 s and 478,126 KiB for the coordinate-only index. The following measurements use the specialized junction-query commands.

For dense, sparse, and junction-set queries, ratios of archive bytes read with versus without the junction-by-shape index are 0.105, 0.129, and 0.105. Wall ratios are 0.308, 0.292, and 0.308. Peak-RSS ratios are 0.189, 0.221, and 0.191. For region queries, the corresponding byte, wall-time, and peak-memory ratios are 1.000, 1.038, and 1.000. Across 96 point-junction queries, median and 95th-percentile source-byte ratios are 0.119 and 0.332. Including collection-index bytes they are 0.175 and 0.528. Aggregate wall ratio is 0.387, a 2.59× speedup.

##### S4.9 Cohort-wide event discovery over the collection

The discovery benchmark uses the same eight SEZ archives and the junction-by-shape collection. `collection find-events` pools the junction catalogues of all archives (1,344,270 junctions), enumerates every cassette whose three components each have support in at least two archives (5,413,686 definitions; 2,976,274 distinct candidate entities), counts exact UMI-class support for every candidate in every archive and cell group, and retains candidates observed in at least four samples and four donors with at least eight informative UMI classes, at least two classes on each side, and at least two classes in each of two required groups (astrocyte/neural stem and mature neuron). Nine cell groups from the transcript-end analysis define the scope, and no genomic coordinates are given as input.

Genome-wide, the scan takes 8.8 s wall (24 threads) and 3.0 GB peak memory, reads the relevant parts of the archive plus 9.6 MB of auxiliary collection data, and performs 324,154,860 exact match attempts and 44,294,758 annotation comparisons. Of the 6.4 s the command itself reports, exact counting takes 3.9 s, candidate enumeration 1.4 s, and annotation classification 0.4 s.

Executed against the coordinate-only collection, the scan takes 8.8 s and reads the same bytes. This is expected, as genome-wide discovery is a scan, and the junction-by-shape routes help targeted panels rather than this workload. Without an annotation filter, 209,570 candidates pass the cohort rule. Requiring incompatibility with every GENCODE v32 transcript retains 205,735, of which 125,005 fail on strand, 76,172 on an exon boundary, 1,116 on overlap, and 3,442 contain a junction absent from the annotation. The last class recurs broadly (2,408 of the 3,442 are seen in all eight donors) and spans 2,843 genes. It includes the 183-nt FNBP1 cassette with 44 informative UMI classes in 44 nuclei across six donors.

The alternative to the `find-events` approach is to run `query events` on each archive and merge. Over the eight archives and 25 chromosomes this takes 23.9 s for 200 runs plus 8.9 s to merge, and yields 552,167 distinct events of which 1,196 pass the same cohort rule. Per-archive discovery can only propose a cassette whose three components are all in that archive's own catalogue. The FNBP1 cassette appears in the output of two archives (D and G), so the merge sees two donors and rejects it, whereas the collection scan counts its support in six. Candidate definition over the pooled catalogue, followed by exact counting everywhere, is what makes recurrence across donors observable for events that are sparse in any one archive.

#### S5 Biological query and discovery analyses

##### S5.1 Splice-event differences between T cells and monocytes

T cells and monocytes are defined from the STARsolo GENCODE v49 matrices: CD3E $> 0$, LYZ $= 0$ for T cells and LYZ $> 3$, CD3E $= 0$ for monocytes. All event coordinates and molecular counts come from the annotation-free archives. Candidates must be present in at least three of four sample catalogues, have at least ten inclusion-only or exclusion-only UMI classes in both groups in every sample, and have a common nonzero direction of usage difference.

Of 20,028 catalogue-recurrent events, 784 satisfy the depth and direction rule; 44 have absolute usage shift at least 0.10 in every dataset. FYB1 cassette inclusion is 5.7–8.8% in T cells and 98.3–99.1% in monocytes, a −0.903 to −0.932 T-minus-monocyte change. CD47 is another high-effect locus. Both had independent RT-PCR validation in primary immune cells[3]. CD47 was also reported by an annotation-free single-cell splicing analysis[4]. Direct aggregation of STAR junction matrices gives the same directions from a second representation of the same alignments. These exploratory comparisons summarize effect sizes and recurrence across biological samples.

##### S5.2 Observed junction components at AKR1A1

AKR1A1 illustrates how event detection depends on observed junction components. The $D_0$ catalogue contains two of the three junctions needed to define the cassette. In $D_4$, all three are observed and yield 110 informative molecules. The two cell groups with at least 20 informative molecules have inclusion fractions of 0.870 and 0.828, a difference of 4.2 percentage points.

##### S5.3 FNBP1 exon inclusion across tissues and cell groups

For this analysis, two additional PBMC archives extend the federation to six samples across three tissues, multiple chemistry generations, and a 20-fold range in cell count. About 120 GB of raw reads occupy 2.8 GB as archives. One junction predicate returns per-sample, per-cell counts across about 94,000 reporting cells in 0.5 s. Although junctions observed in only one sample depend on sequencing depth, 89–97% of each PBMC sample's junction support mass lies on junctions replicated in other samples.

A PBMC-versus-brain-derived comparison identifies 66 recurrent events. MYL6, RPS24, and PPP1R12A differ in directions previously reported by the SpliZ studies[4,5]. An annotated 183-nt FNBP1 cassette has 4.0–13.1% inclusion in four PBMC archives and 94.1–100% in two brain-derived archives. We note that tissue, tumor, cell composition, and nucleus-versus-cell protocol all vary together in this comparison.

We queried an independent normal human brain dataset[6], requiring at least 50 informative UMIs in called nuclei, archive inclusion of at least 0.75, STAR-junction inclusion of at least 0.50, and agreement in direction. Among 10,261 nuclei, the archive gives 72 inclusion-only and two exclusion-only molecules (97.3%). Targeted annotation-free realignment gives identical event totals and its STAR junction matrix gives 61 inclusion-right versus two skip UMIs (96.8%). These measurements corroborate high inclusion in normal brain.

A marker-based classifier assigns nuclei to five broad cell compartments. Each compartment has two marker sets, called submodules:

| compartment | submodule 1 | submodule 2 |
|---|---|---|
| neuronal | RBFOX3, SNAP25, SYT1, SLC17A7, CAMK2A, SATB2 | RBFOX3, SNAP25, SYT1, GAD1, GAD2, SLC6A1 |
| oligodendrocyte/OPC | PLP1, MBP, MOG, MOBP, MAG, CLDN11 | PDGFRA, OLIG1, OLIG2, CSPG4, VCAN, SOX10 |
| astrocyte | AQP4, ALDH1L1, SLC1A2, SLC1A3 | GFAP, GJA1, S100B, SOX9 |
| microglia/immune | PTPRC, TYROBP, LST1, AIF1, FCER1G, CD74 | P2RY12, TMEM119, CX3CR1, C1QA, C1QB, C1QC |
| vascular | CLDN5, EMCN, PECAM1, VWF, KDR, FLT1 | RGS5, PDGFRB, MCAM, COL4A1, COL4A2, CSPG4 |

Table S5: Fixed broad-cell marker modules used for FNBP1 localization. FNBP1 occurs in neither the features nor the normalization denominator.

For each marker, expression is `log1p(10000 × count / non-FNBP1 GEX UMIs)`, standardized across all 10,261 nuclei and clipped to $[-3, 3]$. A submodule score is the mean marker z-score, standardized again across nuclei. The lineage score is the larger submodule score. The highest lineage is assigned only if its selected submodule detects at least two markers, its score is at least 0.50, and its margin over the runner-up is at least 0.35. Two eligible lineages within that margin, or a nucleus above the 99.5th non-FNBP1 library-size percentile with an eligible runner-up, is labeled ambiguous/doublet.

Event-bearing nuclei occur in every broad compartment. Oligodendrocyte/OPC has the largest count, 30, and a rate of 9.36 per 1,000. All 70 informative molecules outside microglia/immune support inclusion. The two skips occur among the four microglia/immune informative molecules, an inclusion proportion of two of four. With so few molecules this rate is poorly determined. Its 95% Wilson score confidence interval for a binomial proportion[7] runs from 0.15 to 0.85. The targeted archive contains 6,584 of 10,261 called nuclei because a nucleus without a local read has no dictionary entry. Detection rates therefore use all called nuclei assigned to each group as the denominator. The marker classifier assigns 3,533 nuclei to the unresolved category and 303 to the ambiguous/doublet category.

#### S5.4 FNBP1 junction co-occurrence in SEZ donors

Across eight adult SEZ donors, all 39 informative FNBP1 cassette molecules support inclusion and are distributed among seven donors. Donor G has the greatest coverage and is the only donor with at least ten informative molecules. Its locus-wide splice graph contains 24 junction-set patterns and 51 strand-specific UMI-class observations. Three reverse-strand patterns each contain two junctions and together account for six UMIs.

Restricting the common graph to edges observed in at least two donors yields 12 junction-set patterns and 86 UMI-class observations across the cohort. Coverage is concentrated in donors D, E, and G, each with at least ten reverse-strand observations; 63 of the 96 donor-by-pattern counts are zero.

#### S5.5 Terminal fragment-boundary redistribution

The local boundary analysis assigns each unique-placement class a transcript-oriented fragment boundary, deduplicates by coordinate, strand, cell, and class, and clusters within 24 bp. A boundary-cluster-by-group likelihood-ratio test with BH correction evaluates 11,166 genes in 14 s on $D_0$. T-cell versus monocyte comparison yields 15% of testable genes above the within-population null's 95th percentile, compared with 5% expected. PTPRC, CD44, and LCP1 lead the result. In an independent $D_1$ analysis with its own markers and null, 26 of 27 $D_0$-significant genes testable in both datasets are significant again, compared with 5.3 expected by chance.

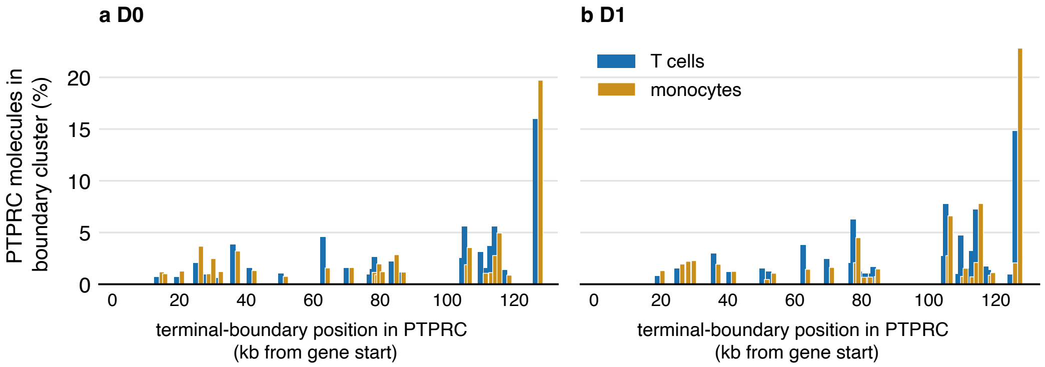


Figure S6: PTPRC terminal fragment-boundary distributions by population, computed independently from two PBMC archives. Fragment boundaries redistribute near a shared terminal region.

For 30 of 35 significant genes, both populations' dominant clusters lie within 200 bp of annotated transcript 3′ ends. Only three of those thirty change the single dominant cluster. The common pattern is therefore redistribution of fragment boundaries near shared termini. Among sites supported by at least 100 molecules, 59.9% of genome-filter-retained sites are within 50 bp of a PolyASite cluster, compared with 11.4% of internal-priming-flagged sites. Agreement with an external site catalogue helps distinguish terminal signals from sequence artifacts that can recur across samples.

#### S5.6 Recovery of genes added in a later annotation

Discovery groups evidence left unassigned by a supplied annotation. The baseline procedure clusters alignment spans outside annotated loci, separately by strand, allowing gaps of up to 1 kb and requiring ten UMI classes. An additional procedure includes molecules overlapping annotated loci when all of their alignments are incompatible with annotated transcripts. Their terminal bases are grouped into windows of at most 1,001 bp. Each proposed locus becomes a single-exon provisional transcript. Replaying this provisional annotation then assigns evidence to transcripts and collapses UMIs.

To measure recovery beyond GENCODE v32, we use 367 genes added after v32, present in v49, and supported by at least 50 STARsolo v49 UMIs in the fixed $D_0$ cell set. Proposed loci are matched one-to-one to these genes by interval overlap, choosing the largest possible set of matches. The baseline procedure recovers 164/367 genes (44.7%).

For the additional overlapping evidence, we compare minimum support levels of 25, 50, 75, and 100 UMIs. We select the level recovering the most genes while proposing at most three times as many loci as the baseline and requiring at least 95% of proposed loci to recur on the same strand in $D_1$. At 75 UMIs, recovery increases to 266/367 genes (72.5%) and 81.5% of their reference UMI mass, with 26,787 proposed loci (2.16× baseline) and 99.0% recurrence.

Recovery is 51/52 for genes outside v32 loci, 52/60 for genes with opposite-strand overlap, and 163/255 for genes with same-strand overlap. Lower support thresholds recover 90.2% and 83.7% of the reference genes while proposing more than three times the baseline number of loci. GENCODE v49 supplies the later-annotation reference. Candidates unmatched to it remain uncharacterized. On a withheld-gene panel, replay of provisional transcripts gives 7.6% median count error, compared with 26% for counts obtained directly from the discovery clusters.

##### S5.7 Reference optimization as replay

The programmatic Pool-style annotation[8] changes 507,335 terminal exon records. Each isoform extends to a shared gene-level boundary, clipped at a neighboring gene and at most 3 kb beyond the gene's outer end. Growth from an internal isoform end can be much longer. Its broad Gene gain therefore includes conversion of intronic intervals into exonic candidates. The span audit found 78,666 expanded gene spans and no contracted spans among 78,691 chromosome/strand/gene combinations.

On the fixed 6,460 brain nuclei, the saved blanket annotation changes Gene counts from 24,153,768 to 54,552,562 (+125.9%), but GeneFull counts from 71,156,828 to 70,576,047 (−0.82%). All exon intervals are preserved or expanded, so the net loss under GeneFull occurs through changed assignment and UMI filtering, not contracted gene spans. Of course, increased counts alone do not establish improved terminal-site inference.

The saved evidence-guided alternative modifies 22,904 genes and excludes unsupported gaps, neighboring-gene corridors, and reference-verified internal priming. On the same nuclei it adds 4,608,988 GeneFull UMIs (+6.48%), compared with 5,261,644 Gene UMIs (+21.78%). Thus 87.6% of the absolute Gene gain remains under intron-inclusive counting, although the larger baseline reduces the percentage gain.

Count gains depend on the barcode scope. Over all raw barcodes, including uncalled droplets, the evidence-guided extension raises Gene totals by 20.15% (28,630,629 to 34,400,997) and GeneFull totals by 6.32% (77,643,934 to 82,548,867). The larger per-nucleus percentages reported above are taken over the 6,460 called nuclei only. On the PBMC dataset the Gene extension raises counts by 16.4% under the blanket extension and 2.3% under the evidence-guided alternative.

#### S6 Terminal-boundary, PolyASite, and tail analyses

##### S6.1 Fragment geometry and analysis design

Fragmented 10x 3′ cDNA boundaries are distributed broadly upstream of RNA cleavage sites. Single-linkage endpoint clusters can also join distinct sites across a terminal region. The mixture model below therefore uses external PolyASite candidates and estimates the distribution of fragment distances from those sites.

We note that the analysis is exploratory, since model choice and selection of NTRK2 used the same eight-donor cohort. Sensitivity to site merging, within-donor label permutations, held-out prediction, and simulations are described below.

##### S6.2 Candidate sites and assay geometry

PolyASite[9] candidates are assigned only to a unique same-strand union of GENCODE terminal exons extended 2 kb downstream. Candidates are merged within 24 bp and rejected when the reference genome has at least 12 A in the downstream 20 bases or an eight-A run within 140 bases.

For endpoint $e$ and candidate $s$, transcript-upstream distance is $d = s - e$ on the forward strand and $d = e - s$ on the reverse. Classes compatible with exactly one candidate estimate a fragment-distance distribution $K(d)$ on $[-50, 2000]$ bp in 10-bp bins. A five-bin moving sum with one pseudocount is normalized. Donor $i$ uses $K_{-i}$ learned from the other seven donors, pooling cell groups during estimation of fragment geometry.

For donor $i$, group $r$, endpoint count $y_{ire}$, and candidate likelihood $L_{es} = K_{-i}(d_{es})$, EM updates

$$z_{ires} = \frac{\theta_{irs} L_{es}}{\sum_h \theta_{irh} L_{eh}},$$

$$N_{irs} = \sum_e y_{ire} z_{ires}, \quad \theta_{irs} = \frac{N_{irs}}{\sum_h N_{irh}}.$$

Initialization is uniform. Iteration stops at maximum $\theta$ change below $10^{-8}$ or 50 iterations.

Sites recur when expected support is at least ten in three donors. Gene usage is the expected-UMI-weighted transcript-oriented site rank. A donor is eligible when both groups have at least 20 expected UMIs. Paired t-test p-values are BH-corrected[10] across genes. Exact sign-flip p-values are also reported. Findings require absolute effect at least 0.10, concordance in six donors, and no leave-one-donor-out sign reversal. A first fit defines recurrence, and a second fit reports support for the common site set in every donor and group, including zeros.

##### S6.3 Predictive and resolution controls

The 24-bp catalogue contains 45,705 recurrent candidates and 35.7M assigned expected UMIs. It tests 2,182 genes and reports 95 under the combined q-value, effect, concordance, and leave-one-out criteria. Held-out prediction improves over a uniform distance model in all donors by 0.418–0.598 bits per unique-candidate UMI. A within-donor label shuffle reports no genes. Ninety-three of 95 primary effects preserve sign at both 12-bp and 48-bp merging choices.

Two-site simulations quantify resolution. At 1,000 UMIs, mean absolute mixture error is 0.054 for sites separated by 50 bp, 0.020 at 250 bp, and 0.013 at 1 kb. Across eligible genes, the median shift toward distal sites is positive in every donor but small (across-donor median 0.013 in distal usage index). The gene-level contrasts below describe larger, locus-specific differences against this small transcriptome-wide shift.

##### S6.4 NTRK2 terminal groups

NTRK2 has mean mature-neuron-minus-Astro/NSC distal-usage effect $+0.282$ ($q = 1.48 \times 10^{-5}$), with all eight donor effects positive. Independent clustering of GENCODE protein-coding transcript ends places four fitted sites beside a proximal GENCODE-Primary group with median 1,431-nt CDS and five beside the distal group containing the MANE Select/full-length transcript, median 2,466 nt. The distal group's pooled share is 20.0% in Astro/NSCs (564 versus 2,260 expected UMIs) and 78.1% in mature neurons (3,146 versus 880). Median paired change is +56.8 percentage points. Exact sign-flip $p = 0.0078$.

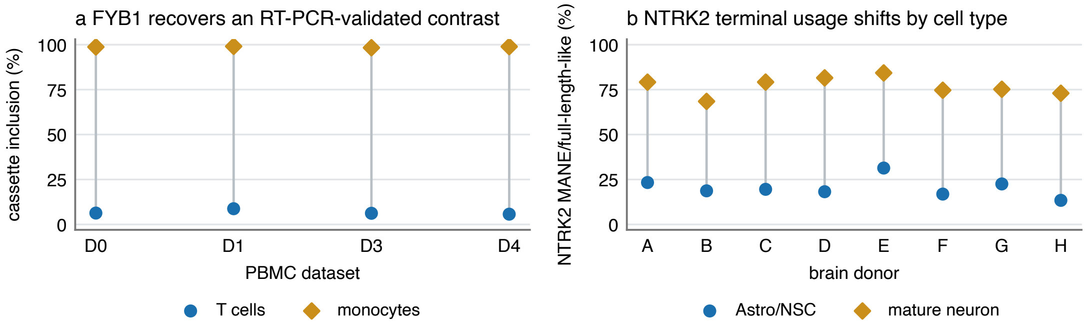


Figure S7: Splice-event and terminal-site examples. The terminal-site panel reports expected usage of groups defined by their proximity to annotated protein-coding transcript ends.

The result agrees with reported predominance of truncated TrkB.T1 in astrocytes and full-length TrkB in neurons[11]. The shorter-coding and MANE/full-length-like labels describe associations between fitted terminal sites and annotated coding ends. The cohort fit takes 28.21 s at 2,262,980 KiB peak RSS, compared with 114.0 s for eight independent donor scans. Processing archives sequentially limits active decoding memory, although retained cell/site evidence still grows with the dataset.

##### S6.5 Non-templated poly(A)-tail evidence

The tail classifier runs during archive construction while alignment-oriented sequence is still available, then discards bases and quality. Under `SoloStrand=Forward`, a forward alignment contributes only its trailing soft clip and is tested for A. A reverse alignment contributes only its leading soft clip and is tested for T. Hard clips do not count. A read is positive exactly when the relevant clip is at least 6 nt, at least 80% of its bases are the oriented tail base, and the transcript-distal homopolymer run is at least 4 nt. The cleavage anchor is the 0-based exclusive aligned end on `+` and the 0-based inclusive aligned start on `-`. Only primary mapped reads with an ACGT UMI and a corrected called-cell barcode under the construction rule in Note S3 are eligible.

Positive reads are deduplicated by `(reference, anchor, strand, corrected barcode, UMI)`, retaining the strongest signal ordered by tail fraction, terminal run, then clip length. For the measurements below, tail evidence is stored in a separate `AIETAIL1` file with a JSON header recording reference names, record count, thresholds, and strand convention, followed by a zstd-compressed stream of fixed 16-byte records:

| field | width | semantics |
|---|---|---|
| reference | 16 bits | index into header reference names |
| anchor | 32 bits | strand-oriented cleavage anchor defined above |
| corrected barcode | 32 bits | 2-bit packed 16-nt barcode |
| UMI | 32 bits | 2-bit packed UMI |
| signal | 16 bits | strand bit; 5 bits each for clip, tail count, and run, saturated at 31 |

Table S6: `AIETAIL1` tail-evidence record, stored separately from the default archive.

A positive read is assigned to an external site when its anchor lies within 25 bp of a PolyASite candidate. The matched control is 500 bp transcript-downstream and is removed when it lies within 100 bp of another external site. Candidate discrimination uses events exclusive to one class and compares clip length with the continuous lexicographic tail-fraction/run score. Across 715,967,790 mapped primary reads from eight donors, 2,932,436 have an eligible terminal soft clip, 144,673 reads pass the classifier, and exact-key deduplication leaves 136,763 positive reads; 53,346 lie near an external candidate. One of those reads also lies in the shifted-control window and is excluded from the exclusive two-class comparison. External candidates therefore have 53,345 positives among 186,470 clipped reads (28.61%), compared with 763/20,386 (3.74%) at shifted controls (odds ratio 10.299). Positivity is depleted at genome-flagged internal-priming sites: 2,085/23,177 (9.00%) versus 51,260/163,293 (31.39%).

The eight compressed tail-evidence files occupy 1,671,772 bytes, or 0.126% of the 1,326,036,691 archive bytes. Only three positive reads occur at NTRK2: one Astro/NSC and two mature-neuron molecules, all in donor G and all at the proximal shorter-coding group.

A UMI class can contain reads with several positive terminal coordinates, including coordinates found on middle reads removed by read reduction. The optional `--terminal-tails` representation extracts these events before reduction and deduplicates by `(cell, UMI class, chromosome, strand, anchor)`. Separate indexed sections store the sparse event lists, with delta-coded record identifiers and anchors and one 16-bit signal word per event.

This representation admits only uniquely mapped primary nonsupplementary reads with explicit `NH=1`. Qualification and selection of the strongest signal use unsaturated counts before the three stored signal counts are saturated at 31. Each deduplicated event stays with the retained record of its strongest supporting read, including when its anchor arose only from a discarded middle read. The stored evidence consists of distinct terminal coordinates and capped signal counts. Tail availability is recorded explicitly in archive metadata.

The 0.126% storage overhead and biological results above apply to the separate `AIETAIL1` files and their primary-read eligibility rule. The integrated option uses the unique-mapping rule described above and is enabled during construction from alignments containing sequence.

### S7 Inference for multi-gene evidence

#### S7.1 Evidence classes and pooled EM

Each UMI class is categorized as single, mixed (unique support for one gene plus alternatives), multi-only, or contradictory. GeneFull ambiguity can arise from overlapping gene spans at one placement as well as alternative genomic placements. GeneFull ignores alternatives on unannotated contigs. In that case the Gene control instead drops the entire class from counting, as the Gene rule does by default. Contradictory classes are excluded. For target class $k$ in cell $c$ with annotation-derived candidate set $C_k$, responsibilities are

$$r_{kg} = \frac{w_{cg}}{\sum_{h \in C_k} w_{ch}}, \quad g \in C_k.$$

Uniform allocation uses $w = 1$. Per-cell EM derives $w$ from one cell.

Pooled EM weights each candidate gene by a sample-wide abundance, $\Pi_g + 10^{-9}$. Each iteration recomputes that abundance as the fixed unambiguous counts plus, for every ambiguous class, the responsibility its candidates receive under the current abundance. The unambiguous counts are the observed part and do not change while the ambiguous contribution is re-estimated from the previous iteration's abundance, so the responsibilities move until they converge. Ten iterations are used.

Only pooled responsibilities for multi-only classes are emitted, as an additive real-valued layer that leaves the base Gene or GeneFull matrix unchanged. The layer counts UMI classes before one-mismatch collapse, so its mass is not a collapsed-UMI total.

##### S7.2 Masked recovery and leakage controls

The evaluation begins with classes whose unique evidence identifies one true gene and then removes that evidence down to the remaining multi-gene candidate set. It scores top-1 accuracy, negative log loss, Brier score[12], and calibration. For an exact responsibility tie, top-1 selects the last gene in the stored candidate order. Every estimator uses the same deterministic convention. Candidate genes are excluded from any replay features used to construct evaluation groups, preventing hidden-label leakage through clustering. Uniform, per-cell, pooled, and group-aware estimators use identical targets.

All four primary panels were rerun with 20% masking, seed 7, ten iterations, and blend weight 20. The original called barcodes restrict scoring. All archive barcodes contribute to the fitted pooled prior after masking. The group map is constant and supplies only the scoring population for the four base modes below. A mixed class is eligible when exactly one gene has unique support. Its label must survive in the remaining multi-gene candidates to enter conditional accuracy. Counts below are UMI classes before one-mismatch collapse, not reads, archive records, or collapsed matrix UMIs.

| dataset / model | masked | evaluable | uniform | per-cell | pooled |
|---|---|---|---|---|---|
| $D_0$ Gene | 171,089 | 166,755 | 53.3% | 96.6% | 98.2% |
| $D_1$ Gene | 880,529 | 865,224 | 51.0% | 96.6% | 98.3% |
| $D_2$ Gene | 253,471 | 245,734 | 57.9% | 92.6% | 95.7% |
| $D_{2'}$ GeneFull | 231,351 | 156,884 | 43.2% | 68.9% | 74.9% |

Table S7: Masked recovery on fixed called barcodes. Percentages divide correct top-1 assignments by evaluable classes. Masked minus evaluable counts labels lost from the remaining candidate set.

For brain GeneFull, 74,467/231,351 masked classes (32.2%) lose their label. Pooled recovery is 74.9% among the remaining 156,884, versus 68.9% per cell and 76.1% for blend. Counting lost labels as failures instead gives 50.8% pooled recovery among all masked classes. Seeds 17 and 29 give 74.5% and 74.7% conditional pooled recovery. Only 80,412/156,884 targets (51.3%) have maximum pooled responsibility at least 0.9; 74,650/80,412 (92.8%) of those are correct. This differs from $D_0$, where 161,288/166,755 (96.7%) fall in that bin and 160,141/161,288 (99.3%) are correct. Responsibility thresholds therefore need dataset-specific calibration, or a more sophisticated model to determine them.

On the same brain nuclei, Gene gives 72.7% per-cell and 88.5% pooled recovery on 70,278 evaluable classes, with 2,393/72,671 (3.3%) labels lost. Gene and GeneFull change the eligible population as well as candidates, and so this is not a paired comparison on identical truth classes. Scoring every archive barcode instead gives 68.6%/90.9% per-cell/pooled under Gene (100,186 evaluable classes), and 66.7%/76.8% under GeneFull (193,141). Of course, none of these unique-evidence labels establishes independent biological accuracy.

The separate $D_0$ emission experiment adds inferred mass equal to 6.3% of single-gene-supported UMI classes before one-mismatch collapse, 85% at responsibility above 0.8.

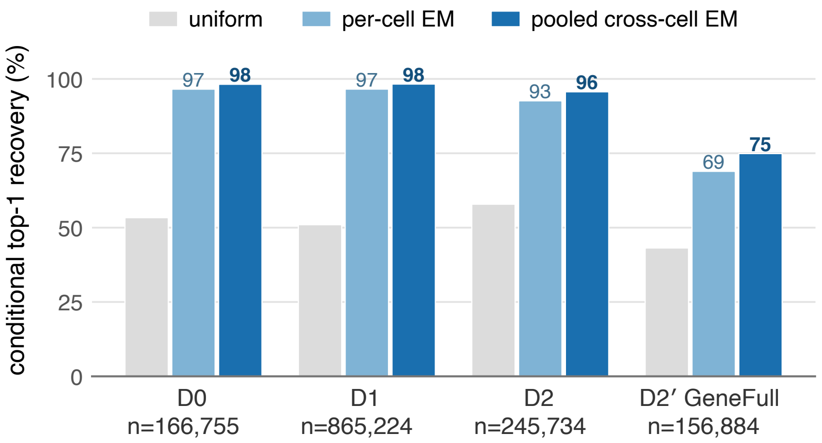


Figure S8: Conditional masked recovery on fixed called barcodes. Brain nuclei use GeneFull, other datasets Gene; $n$ counts evaluable UMI classes. Sample pooling is the default emitted estimator. Group-aware partial pooling is evaluated separately.

External corroboration uses alevin-fry, which has access to read sequence that the archive discards. Among the 254 genes receiving most recovered mass, unique-only counts differ from alevin-fry by median 2.5×. Adding the layer reduces this to 11%, increases log-correlation from 0.79 to 0.87, and moves 79% of genes closer. Recovered-mass fractions agree between two PBMC datasets at Pearson $r = 0.992$ with median absolute difference 0.004 across 2,823 genes.

##### S7.3 Candidate-normalized partial pooling

Partial pooling combines evidence from a cell, its expression-defined group, and the whole sample. Each distribution is normalized over the target class's candidate genes, so the specified pseudocount mass applies to that candidate set. Let $\pi_{cg}$ be the current abundance of gene $g$ in cell $c$, $\Pi_{hg} = \sum_{c \in h} \pi_{cg}$ the abundance in its group $h$, and $\Pi_g = \sum_c \pi_{cg}$ the sample abundance. For candidate set $C$, define

$$p_0(g|C) = \frac{\Pi_g + \varepsilon}{\sum_{j \in C} (\Pi_j + \varepsilon)},$$

$$p_{c(g|C)} = \frac{\pi_{cg}}{\sum_{j \in C} \pi_{cj}},$$

$$n_{hg,-c} = \max(\Pi_{hg} - \pi_{cg}, 0),$$

$$p_{h(g|C)} = \frac{n_{hg,-c} + \kappa p_0(g|C)}{\sum_{j \in C} n_{hj,-c} + \kappa}.$$

Here $\varepsilon = 10^{-9}$ supplies a small positive sample weight and $\kappa = 80$ is the sample-derived pseudocount mass added to the group. At 80 observed group counts among the candidate genes, group evidence and the sample prior contribute equal mass. When the cell denominator is zero, $p_c = p_0$. The group estimate excludes the target cell. The fixed convex combination is

$$q_{\text{conv}}(g|C) = 0.20 p_{c(g|C)} + 0.60 p_{h(g|C)} + 0.20 p_0(g|C).$$

The weights and $\kappa$ were selected by minimizing mean negative log loss in masked recovery on $D_0$, using masking seeds 7, 17, and 29. Cell weights were 0, 0.05, 0.10, or 0.20. Group weights were 0, 0.25, 0.50, 0.75, or 1 times the weight remaining after the cell contribution. The sample received the remaining weight. For positive group weights, $\kappa$ was 0, 5, 20, or 80. Zero-group settings used $\kappa = 20$. Ties favored lower group weight, then lower cell weight, then lower $\kappa$. The selected combination assigns 20% to the cell, 60% to its group, and 20% to the sample, with $\kappa = 80$. These are empirically selected smoothing parameters for the masked-recovery task. While it is outside the scope for the current manuscript, we suspect a more sophisticated model may allow a more principled way to learn these weights that adjusts to the datasets being analyzed.

If a cell lacks a group assignment, the group weight is transferred to the sample term, giving $0.20 p_c + 0.80 p_0$. Empty cell or group evidence therefore has a defined sample fallback, and all distributions have support only on $C$.

A second estimator treats the group and cell distributions as successive posterior-mean proxies. With fixed group and cell prior masses 80 and 64,

$$p_{\text{group}}(g|C) = \frac{n_{hg,-c} + 80p_0(g|C)}{\sum_{j\in C} n_{hj,-c} + 80},$$

$$p_{\text{proxy}}(g|C) = \frac{\pi_{cg} + 64p_{\text{group}}(g|C)}{\sum_{j\in C} \pi_{cj} + 64}.$$

These equations shrink cell estimates toward their group and group estimates toward the sample. The prior masses have units of candidate-gene counts: 64 makes the cell evidence and group-derived prior equally weighted when the cell has 64 counts across its candidate genes. The group prior has the analogous interpretation at 80 counts. Both masses were selected on $D_0$ by minimizing mean negative log loss over the same three masking seeds. The search combined cell prior masses of 1, 4, 16, 64, and 256 with group prior masses of 1, 5, 20, 80, and 320. Ties favored the lower cell prior, then the lower group prior. The selected values, 64 and 80, were held fixed in subsequent comparisons, including the held-out $D_3$ analysis below.

##### S7.4 Depth-dependent combination

Cell evidence should matter more when the cell contains appreciable unique mass for the target's candidate genes. Let $d = \sum_{g\in C} u_{cg}$, where $u$ is the initial unambiguous count table after masking, before any fitted responsibility is added. Thus $d$ is fixed before optimization and is independent of which estimator is being compared. The depth weight

$$s(d) = \frac{d^P}{d^P + D^P}, \quad D = 8, \quad P = 8$$

is bounded and monotone in candidate-relevant depth. The evaluated estimator is

$$q_{\text{hybrid}}(g|C) = (1 - s(d))q_{\text{conv}}(g|C) + s(d)p_{\text{proxy}}(g|C).$$

Both component estimators are recomputed from one evolving abundance estimate. In each of ten deterministic iterations, every target is evaluated with $q_{\text{hybrid}}$ from the current state. The next cell, group, and sample state is reset to the unambiguous table and accumulates those same hybrid responsibilities at all three scales. Recomputing both components from this shared state keeps their latent-count estimates consistent.

The depth weight changes only the relative contribution of two candidate-supported distributions. Responsibilities are nonnegative and sum to one over $C$. As cell depth increases, $s(d)$ shifts weight toward the cell-adaptive estimator.

##### S7.5 Leakage-controlled group construction

Groups are constructed from Gravlax Gene replay against GENCODE v49. The union of candidate genes across all targets in the three masked-recovery replicates is removed before normalization, feature selection, principal components, or clustering. Fixed STARsolo called-cell barcodes define the cell universe. Genes detected in at least ten cells are retained. Each cell is normalized to 10,000 counts and transformed by `log1p`. At most 2,000 highly variable genes are standardized before PCA.

The grid uses 20, 30, 40, or 50 PCs, Leiden resolution 0.20–1.00 in increments of 0.05, and seeds 0–19. At each grid point the seed with maximum mean adjusted Rand index (ARI) to the other nineteen partitions is the medoid. A point is eligible only when it yields 3–12 groups, every group has at least 20 cells, and the median of the 190 pairwise seed ARIs is at least 0.90. Selection maximizes median ARI, then prefers fewer PCs, resolution closest to 0.50, and lower resolution. A deterministic size-preserving permutation of the final labels supplies the negative group control.

On held-out $D_3$, candidate-excluded replay yields 11 groups among 11,644 cells with 50 PCs and resolution 0.30. Median pairwise seed ARI is 0.986 and the smallest group has 24 cells. Across masking seeds 7, 17, and 29, real groups improve target-count-weighted mean negative log loss by 1.074% relative to the size-preserving shuffled groups. Expression-defined groups therefore provide predictive information about gene assignment after candidate genes have been excluded from clustering.

##### S7.6 Parameter selection and uncertainty

Depth parameters were selected on $D_0$ and $D_4$ using $D \in \{2, 4, 8, 16, 32, 64, 128, 256\}$ and $P \in \{0.5, 1, 2, 4, 8\}$, with masking seeds 7, 17, and 29 and the real stable groups above. A pair was eligible only if, separately on both datasets, its target-count-weighted mean negative log loss was lower than both pooled EM and the proxy estimator, its Brier score was no greater than either, and its top-1 accuracy was no more than 0.10 percentage point below the better estimator. Among the 24 eligible pairs, selection maximizes the minimum relative Brier improvement over the proxy across $D_0$ and $D_4$. Ties use lower joint negative log loss, then lower $P$ and lower $D$. This rule selects $D = 8, P = 8$, which is carried unchanged to the held-out $D_3$ evaluation.

Loss comparisons are paired by target. Within a masking seed, standard errors cluster paired loss differences by target cell and form normal 95% intervals. Across seeds, target counts weight the estimates. The reported standard error is the same weighted sum of seed-level clustered standard errors, equivalent to a conservative perfect-positive-correlation assumption because seeds reuse cells and have overlapping targets.

Relative to sample-pooled EM on $D_3$, the hybrid changes negative log loss by −0.00247187 (95% CI [−0.00359083,-0.00135291]) and Brier score by −0.00168846 ([−0.00252929,-0.000847637]). Relative to the proxy estimator, the changes are −0.000208191 ([−0.000244221,-0.000172162]) and −0.0000905761 ([−0.000109928,-0.0000712242]). Top-1 accuracy improves by 0.098 percentage point over pooled EM and 0.005 point over the proxy.

##### S7.7 Memory use

Processing groups of cells individually from temporary files reduces the active memory requirement. On the $D_3$ shuffled-control run with masking seed 7, this approach reduces peak RSS from 7,653,508 to 3,846,008 KiB and completes in 35.86 s. Temporary storage, of course, still grows with input size.

While the hybrid gives the best masked-recovery results among the tested estimators, sample-pooled EM remains the default emitted layer, because its downstream consequences have been evaluated and the hybrid's have not.

#### S8 Datasets, software, and reproducibility

##### S8.1 Evaluation datasets

| ID | material | reads | cells | role |
|---|---|---|---|---|
| $D_0$ | human PBMC 1k, 10x 3′ | 66.6M | 1,225 | primary replay/query |
| $D_1$ | human PBMC 5k, 10x 3′ | 383.9M | 5,038 | scale/replication |
| $D_2$ | human glioblastoma, 10x 3′ | 250.7M | 5,573 | tissue generalization |
| $D_{2'}$ | human brain nuclei, 10x 3′ | 263.4M | 6,460 | sparse/deep-intron |
| $D_3$ | human PBMC 10k v3 | 638.9M | 11,644 | held-out grouped EM |
| $D_4$ | human PBMC LT Chromium X | 75.7M | 500 | EM development/federation |
| $D_5$ | mouse splenocyte, 10x 5′ v2 | 33.7M | 1,129 | protocol/species |

Table S8: Named evaluation datasets. Cell counts are the fixed called-cell sets used by the corresponding analyses.

The following public catalogue identifiers specify datasets $D_0$–$D_5$ and the independent normal-brain dataset used for the FNBP1 confirmation.

| ID | public identity or accession | provider |
|---|---|---|
| $D_0$ | `pbmc_1k_v3` | 10x Genomics |
| $D_1$ | `5k_pbmc_v3` | 10x Genomics |
| $D_2$ | `Parent_SC3v3_Human_Glioblastoma` | 10x Genomics |
| $D_{2'}$ | `Brain_3p` (Cell Ranger 6.0.0 single-nucleus brain) | 10x Genomics |
| $D_3$ | `pbmc_10k_v3` | 10x Genomics |
| $D_4$ | `500_PBMC_3p_LT_Chromium_X` | 10x Genomics |
| $D_5$ | `sc5p_v2_mm_c57bl6_splenocyte_1k` | 10x Genomics |
| brain conf. | `10k Human Brain Nuclei, Chromium GEM-X Epi Multiome`[6] | 10x Genomics |

Table S9: Public dataset identifiers. The final row is the independent normal-brain dataset used only for the prospective FNBP1 confirmation (Supplementary Note S5). Its gene-expression library was used. The exact $D_5$ FASTQ archive is linked here. These datasets are hosted by 10x Genomics under the identifiers shown.

The adult human subependymal-zone cohort is GSE234790/PRJNA983239 and comprises eight donors, 36,626 published-QC nuclei, and 977.3M read pairs[13]. Human analyses use GRCh38 with GENCODE[14] v32, v48, or v49. Mouse uses GRCm39 with GENCODE M39. Unless stated otherwise, commands use 24 aggregate threads.

##### S8.2 Genome and annotation identity

Every archive records reference names and lengths. Sequence-consulting commands require per-contig BLAKE3 signatures and fail on a mismatched FASTA. Compiled annotations are deterministic and checksummed. Collection federation requires matching chromosome digests and genome signatures.

##### S8.3 Compute and software environment

Measurements run on a shared Red Hat Enterprise Linux 8 host with two AMD EPYC 9575F processors, 256 logical CPUs, and 1.5 TiB RAM. Project jobs are capped at 24–32 threads. Paper runtime comparisons use 24 aggregate threads unless a different budget is stated. The alignment and comparison environment uses STAR 2.7.11b[15], samtools 1.23.1[16], and gffread 0.12.9[17].

##### S8.4 Timing and memory measurement

Warm timings were taken without explicit cache eviction on a shared host. Cold-cache timings for specific inputs are reported separately. Runtime records include inputs, annotation, cell groups, thread allocation, wall time, and peak resident set size. Logical file sizes are used throughout.

Collection-query measurements record bytes read from source metadata, source records, and collection-index bytes separately. A query for an absent coordinate reads source identity metadata but can avoid molecule payloads.

Elapsed time includes reading the location manifest. Byte counters measure reads from archives and collection indexes.

##### S8.5 Software and archived workflows

Source and executable workflows are available from COMBINE-lab/gravlax and COMBINE-lab/gravlax-paper-scripts. The format specification provides encoding details. Result bundles record the software revision, input and reference checksums, command-line parameters, annotation, cell-group definitions, and thread budget for each analysis.
